# Understanding the effect of temperature on Bluetongue disease risk in livestock

**DOI:** 10.1101/759860

**Authors:** Fadoua El Moustaid, Zorian Thronton, Hani Slamani, Sadie J. Ryan, Leah R. Johnson

**Author notes:** Corresponding author: Department of Biological Sciences, 2119 Derring Hall, Virginia Tech, Blacksburg, VA 24061 USA.

## Abstract

The transmission of vector-borne diseases is governed by complex factors including pathogen characteristics, vector-host interactions, and environmental conditions. Temperature is a major driver for many vector-borne diseases including Bluetongue viral (BTV) disease, a midge-borne febrile disease of ruminants, notably livestock, whose etiology ranges from mild or asymptomatic to rapidly fatal, thus threatening animal agriculture and the economy of affected countries. Using modeling tools, we seek to predict where transmission can occur based on suitable temperatures for BTV. We fit thermal performance curves to temperature sensitive midge life history traits, using a Bayesian approach. Then, we incorporated these into a new formula for the disease basic reproductive number, *R*_0_, to include trait responses, for two species of key midge vectors, *Culicoides sonorensis* and *Culicoides variipennis*. Our results show that outbreaks of BTV are more likely between 15°C and 33°C with predicted peak transmission at 26°C. The greatest uncertainty in *R*_0_ is associated with the uncertainty in: mortality and fecundity of midges near optimal temperature for transmission; midges’ probability of becoming infectious post infection at the lower edge of the thermal range; and the biting rate together with vector competence at the higher edge of the thermal range. We compare our *R*_0_ to two other *R*_0_ formulations and show that incorporating thermal curves into all three leads to similar BTV risk predictions. To demonstrate the utility of this model approach, we created global suitability maps indicating the areas at high and long-term risk of BTV transmission, to assess risk, and anticipate potential locations of establishment.

## 1 Introduction

With ongoing climate change, it is critical that we understand how temperature influences the dynamics of emerging diseases. Vector-borne diseases are highly sensitive to climate factors, particularly temperature [1, 2, 3]. Bluetongue virus (BTV) is no exception. The biting midges of the *Culicoides* family responsible for transmitting BTV and many other arboviruses are highly sensitive to changes in temperature [4, 5], and thus so is BTV transmission [6, 7]. More than 1,400 species of *Culicoides* have been classified, globally, but fewer than 30 have been identified as competent vectors for BTV transmission [8].

BTV was first detected among merino wool sheep in South Africa in 1905, and since then, the disease has been found on every continent but Antarctica [9]. In recent years, outbreaks have occurred in North and Central Europe, parts of Asia, and Western North Africa [10]. Though the cause of the recent appearance of BTV in some of these regions (especially Northern Europe) is still unknown, it is believed that climate change is the main driver, as the increase in temperature of certain locations makes them suitable for midges to survive, and therefore transmit diseases [11].

BTV can infect most species of domestic and wild ruminants, including sheep, goats, and cattle [12]. Sheep are the most susceptible to the disease and exhibit the highest morbidity and mortality post infection [11, 13]. In the majority of strains of BTV’s 27 serotypes, infected animals rarely show any clinical signs [14]. The severity of infection, and presence of clinical signs depend on the serotype, and the severity of infection can range from rapid fatality to quick recovery. Common outward clinical signs include a blue tongue, fever, and excessive salivation [11].

Since clinical signs are rarely exhibited by infected animals, in many cases BTV goes without detection. Unofortunately, undetected cases can still result in mortality, and BTV vaccine development is in its infancy [15]. An effective polyvalent vaccine to immunize against more than one strain of BTV has yet to be developed [16]. Further, existing attenuated viral vaccines pose significant health risks to livestock, such as reduced milk production in lactating sheep, abortion, early embryonic death, and teratogenesis in pregnant females [17]. With the absence of an effective polyvalent vaccine, and the potential risks and costs of the available vaccines, the impact of BTV on the U.S. beef industry can be significant. This cost was estimated at $95 billion dollars in 2014 [16]. The mandatory testing of animals and losses in foreign markets add to the economic impact of BTV on the U.S. livestock industry, underscoring the need to assess the risk from it. The impact of BTV extends far beyond the U.S.; the spread of the disease to areas previously believed to not be at risk, such as North and Central Europe, regions of Asia, and Western North America, is concerning [13].

Although climate change is believed to be behind this unprecedented spread of BTV to new areas, more research is needed to verify this hypothesis. Some of the cases of BTV-8 in Europe, specifically in France, have exceeded expectations and survived cold winters [18]. With such crucial components such as the potential for geographic spread, and virus survival, still unexplored, it is essential to study BTV further, to uncover the intricacies that dictate its impact.

BTV cannot survive on its own in the environment. As shown in Figure 1, BTV transmission involves host-vector interactions, host-virus interactions, vector-virus interactions, and the effect of the environment. Once BTV is transmitted to a vector or host, its survival is most notably dependent on temperature; while BTV can survive for years at some temperatures, it can be inactivated in a matter of hours within a host at other temperatures [13].

**Figure 1:**
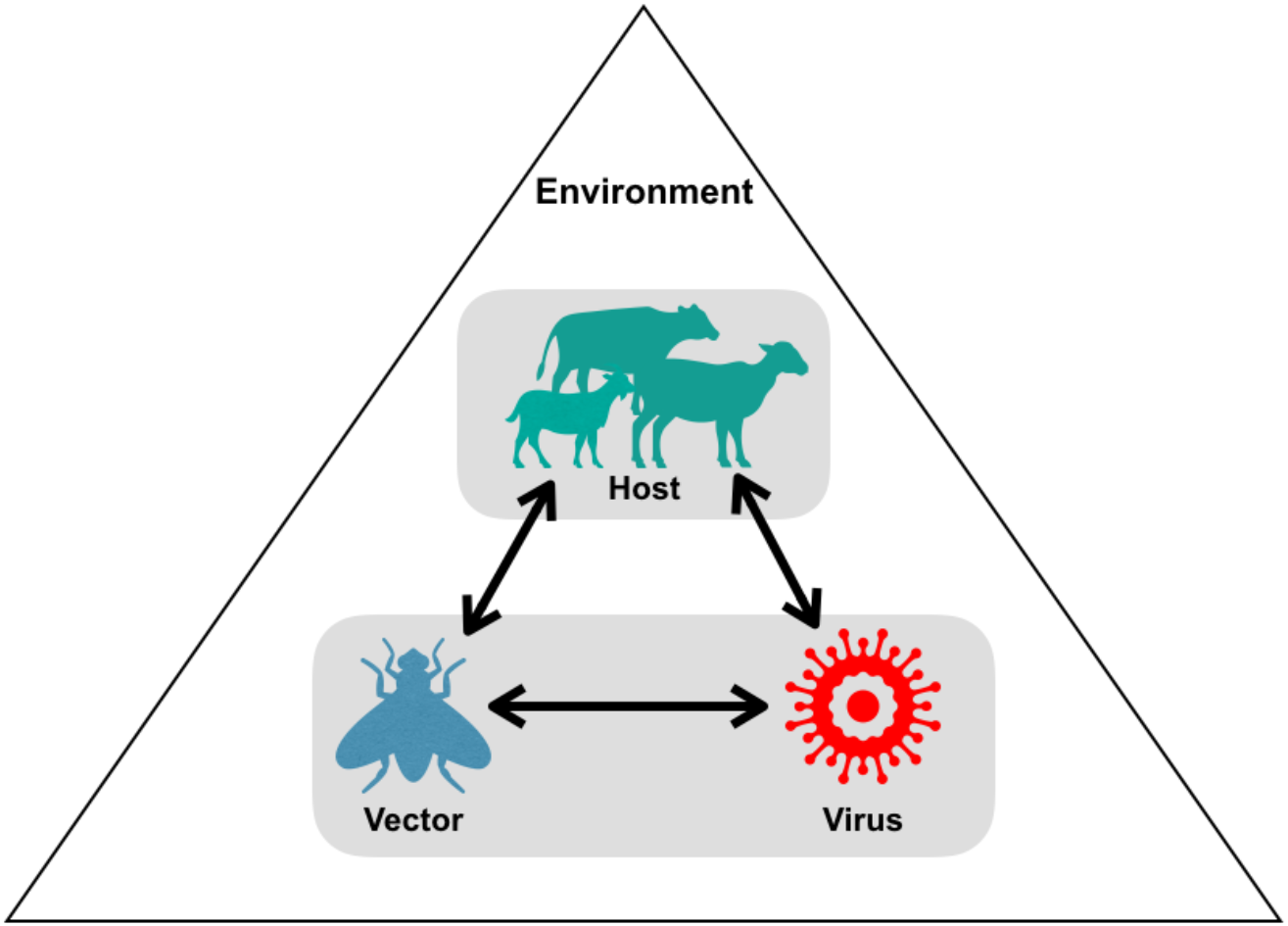
Bluetongue virus transmission diagram: the mechanisms underlying the transmission of bluetongue virus include, host-vector interactions, host-pathogen interactions, vector-pathogen interactions as well as the environmental effect on all interactions.

Due to the complexity involved in determining BTV and other VBD transmission, scientists often use mathematical modeling to understand the transmission process of these diseases [19, 20, 21]. The classical Ross-MacDonald model of VBDs and similar models allow us to calculate the corresponding basic reproductive ratio *R*_0_ of the disease [22, 23]. This summary quantity is widely used to estimate how contagious the disease is and whether an outbreak can occur. When *R*_0_ > 1, the disease is likely to spread, leading to an outbreak, while *R*_0_ < 1, indicates that the disease is likely to die out.

Here we are interested in answering the following questions: (1) How does *R*_0_, and thus risk of transmission of BTV, vary with temperature? (2) Do different model assumptions lead to different values of *R*_0_? and (3) Which traits contribute the most to variation in *R*_0_ estimates? To answer these questions, we take the following three-step approach used previously for VBDs such as malaria [1, 2]. First, we derive a new *R*_0_ for BTV, and we use Bayesian inference to fit lab data for temperature-sensitive midge life-history traits, to estimate the thermal responses of these traits. Next, we compare our *R*_0_ to two previously derived forms where the midge density, *V*, is constant to ascertain if this generates major differences in outbreak prediction. We then contrast this with a temperature-sensitive midge density, *V*(*T*), expressed in terms of midge traits.

We compare results for all three forms of *R*_0_, and identify which parameters lead to high variation in *R*_0_, using uncertainty analyses. This can indicate that either further data collection is needed to refine estimates, or that certain parameters have greater impacts on BTV disease transmission at different temperatures. Furthermore, understanding which temperature range results in *R*_0_ > 1, for given levels of other fixed parameters in our model, could inform prevention and control strategies.

## 2 Methods

### 2.1 Derivation of *R*_0_

To predict the outbreak potential of BTV, several forms of the basic reproductive number *R*_0_ have been developed [19, 20, 21]. The classical reproductive ratio for a generic VBD [1, 24] is given by

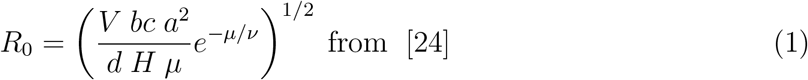

where *V* is midge population density; *bc* is vector competence (the product of the probability that a midge can transmit the infection to an uninfected host, *b*, and the probability that a midge gets infected when biting an infected host, *c*); *a* is the per-midge biting rate; *μ* is the adult midge mortality rate; *ν* is the pathogen development rate (*ν* = 1/*EIP* with *EIP* the extrinsic incubation period); *H* is host density; and *d* is infected host recovery rate. The model used to derive this version of *R*_0_ is a system of delay differential equations that assumes no exposed class and that susceptible midges move to the infected class shortly after contact with an infected host. A similar scenario can be described using a system of ordinary differential equations while expressing the delay between the contact with infected host and midges becoming infectious in terms of an exposed class. In this case, the reproductive number for the midge-borne viral disease (BTV) can be expressed as,

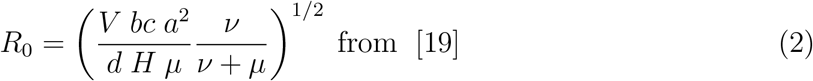

This version of *R*_0_ is a reduced version from a model that uses multiple types of host and multiple types of midge species as in [19, 20].

Figure 2 shows a schematic representation of our BTV transmission model (Equations (S1.1)-(S1.8) in Appendix S1) which considers a single host population split into susceptible individuals that are vulnerable to BTV disease (*S*), infected individuals that have acquired infection (*I*), and individuals who have recovered from the disease (*R*). In addition, we consider a vector population containing susceptible midges (*S_v_*), three levels of exposed individuals (*E_v_*), and an infected class of midges (*I_v_*). The exposed classes in the model represent the extrinsic incubation period that midges undergo before becoming able to transmit infection. To calculate the basic reproductive number *R*_0_, we use a next generation matrix method described in [25, 26], which leads to the following *R*_0_ equation:

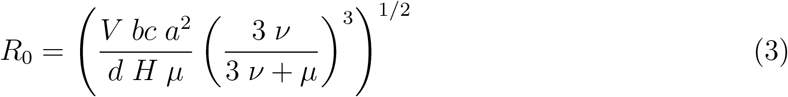

all parameters in our *R*_0_, except *d* and *H*, are assumed to be temperature dependent. The term 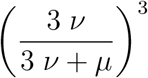 in our *R*_0_ represents the number of midges that survive the extrinsic incubation period, leading to a slight difference between the three *R*_0_ forms. We can represent all three formulas of *R*_0_ with a simple equation given by:

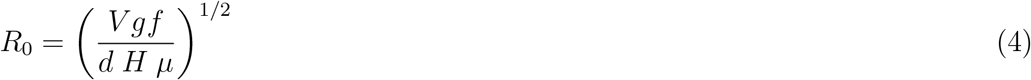

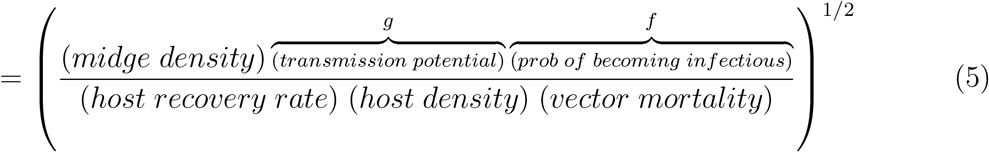

where the expression for *V* and *g* are the same for all three versions and are given by:

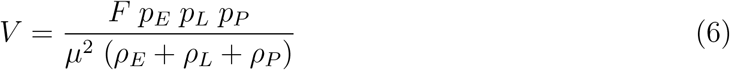

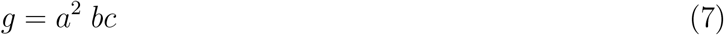

where *F* is eggs per female per day, *p_E_, p_L_*, and *p_P_* are survival probabilities for eggs, larvae, and pupae; *ρ_E_, ρ_L_*, and *ρ_P_* are development times for eggs, larvae and pupae, respectively; *μ* is adult midge mortality; *a* is midge biting rate and *bc* is midge competence. The difference between the three *R*_0_ formulas lies in the functional form, *f*, representing the probability of midges surviving to become infectious post infection. Table 1 summarizes the functional forms for each of the three models considered.

**Figure 2:**
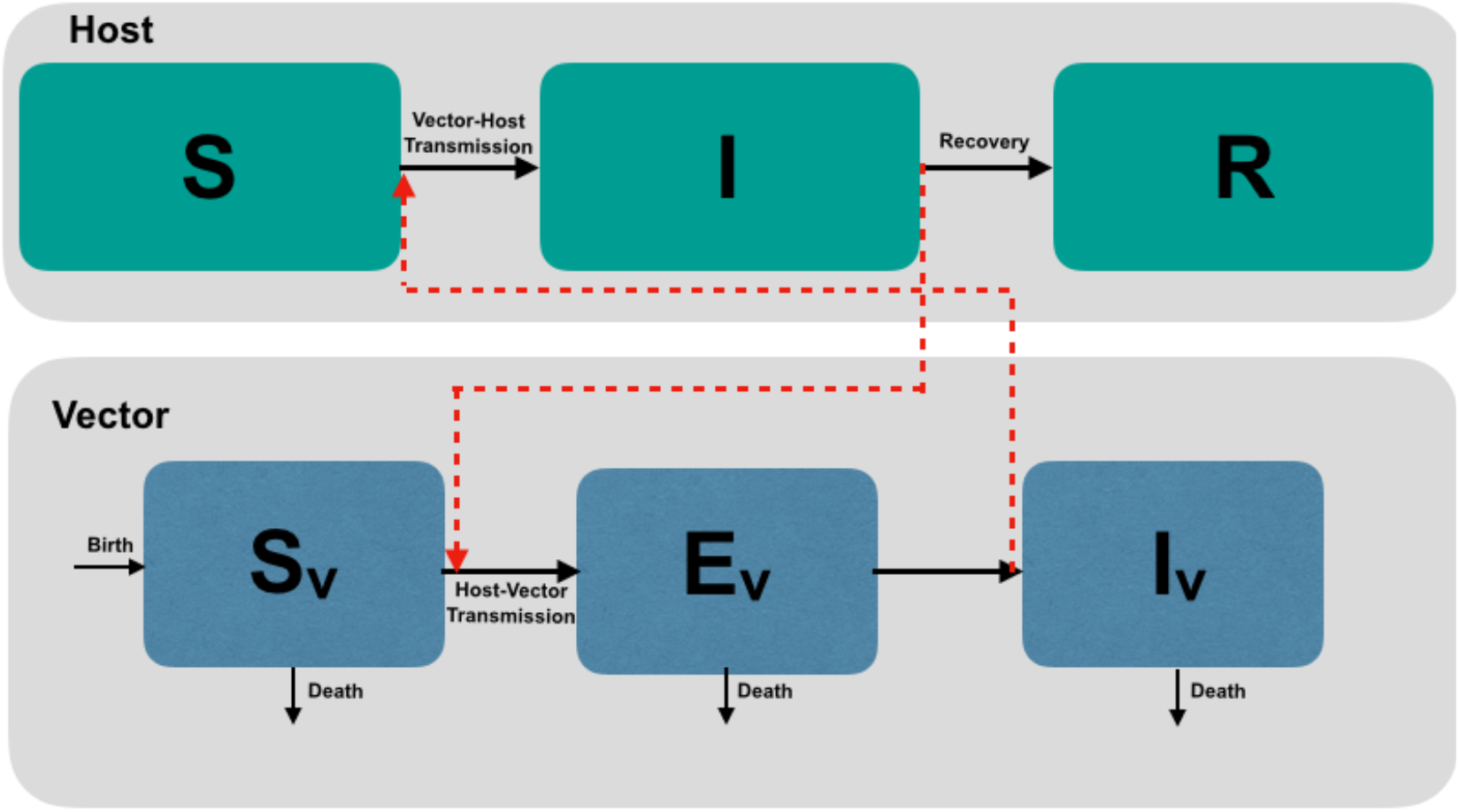
A schematic illustration of BTV transmission. The host population is composed of three classes: susceptible (*S*), infected (*I*), and recovered (*R*). The midge population is composed of a susceptible class (*S_v_*), three exposed classes (*E_v_*), and an infected class (*I_v_*). Black arrows show movement between classes and red arrows indicate contact potentially leading to transmission.

**Table 1:**
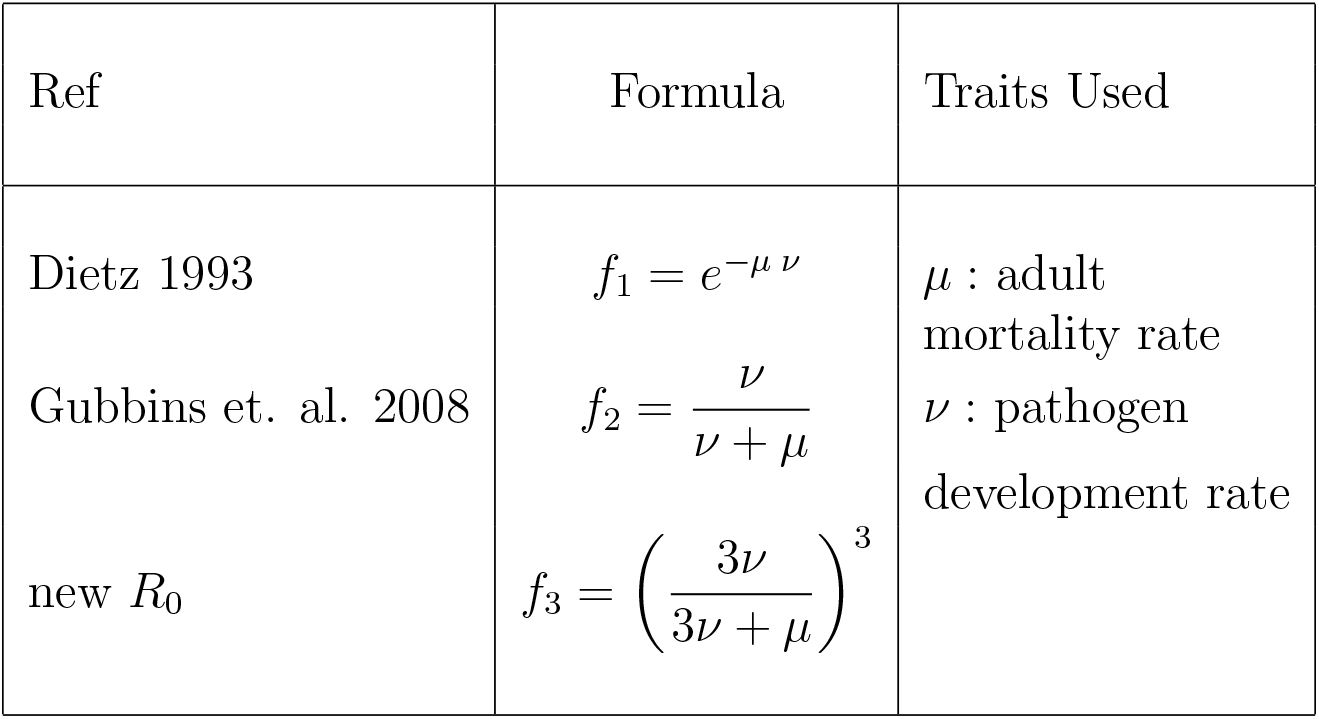
Functional forms used in *R*_0_ formulas and the parameters involved.

Figure 3 shows all three functional forms as we fix the parasite development rate *ν* and vary the midge mortality rate *μ*. We use all three forms in our analysis while comparing the constant vector density case *V* to temperature-sensitive density *V*(*T*).

**Figure 3:**
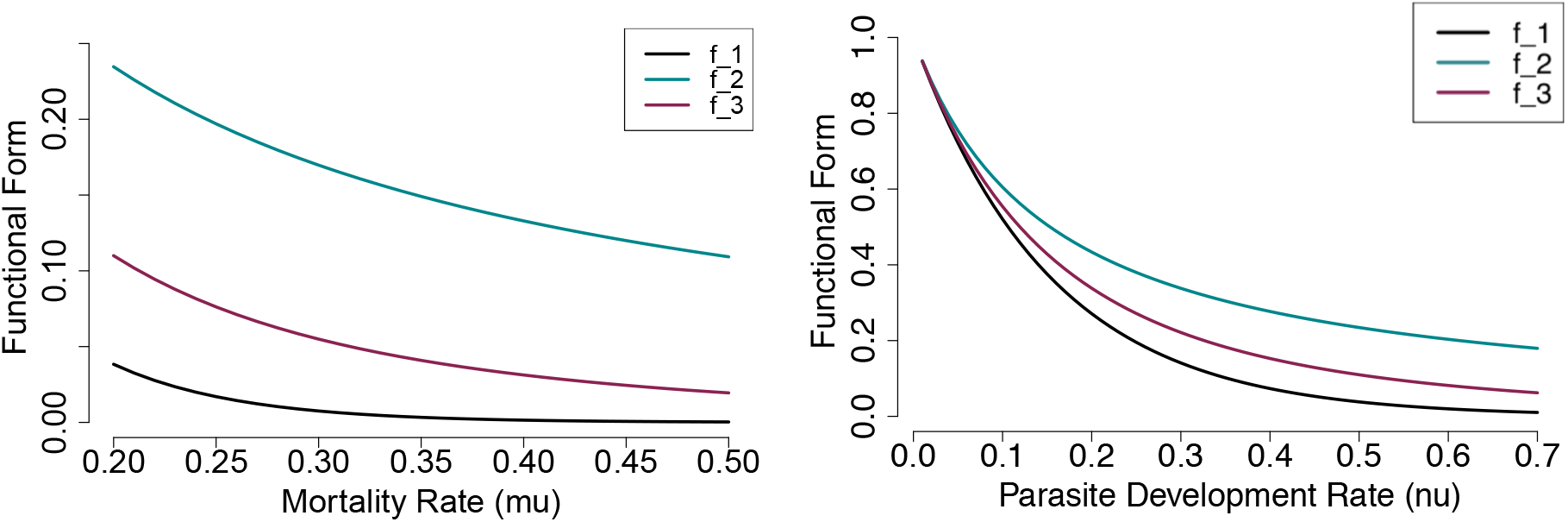
(Left) Functional forms *f* versus midge mortality rate *μ* with a fixed *ν* = mean(*ν*(*T*)) = 0.061. (Right) Functional forms *f* versus pathogen development rate *ν* with a fixed *μ* = mean(*μ*(*T*)) = 0.15

### 2.2 Bayesian fitting of temperature-sensitive traits

As ectotherms, midges are sensitive to temperature. The thermal performance for these temperature-dependent traits is generally hump-shaped, starting at zero at a given minimum temperature, then increasing to a peak value as temperature increases, then sharply dropping to a lower value at a maximum temperature [27, 28].

Here, we collected trait data corresponding to two midge species from the family *Culicoides*, namely, *Culicoides sonorensis* and *Culicoides variipennis*, both found the US [29]. The data collection method consisted of synthesizing data from published literature, via assembling data from tables, and digitizing data points from graph; details on data fitting for each trait is provided in Appendix S1. In searching for the data we would use for this study, we were particularly interested in controlled laboratory experiments on midge trait variation at constant temperatures, ideally with three or more data points. For digitization, we used the free software `P`lotDigitizer [30].

We used the temperature-dependent trait fits in all three *R*_0_ formulas for comparison. Following a method first introduced by Mordecai et al. in 2013 [1], we fit unimodal curves to temperature-sensitive traits. For the unimodal curves, we chose between a Brière (Equation (8)) or a quadratic formula (Equation (9)) depending on the trend seen in data.

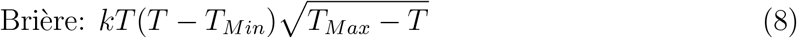

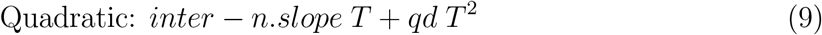

where the constants *k, T_Min_, T_Max_, inter, n.slope*, and *qd* are the result of fitting trait data, and each trait requires specific values for the equation to fit the data. For more information on the values, see Appendix S1.

Similarly to Johnson et al., [2], we used a Bayesian approach for our fitting method. For each continuous trait, we choose a normal distribution as our likelihood that represents the data. We truncated the likelihood distributions at zero to avoid negative values for all thermal curves, and at one for fitting probabilities. We chose priors for each of the fitting form parameters to assure parameters have the correct sign and range. When fitting probabilities, we often switched our likelihood to a binomial distribution.

We use a Markov Chain Monte Carlo (MCMC) sampling in `JAGS/rjags` to fit our models [31]. For each fit, we run five MCMC chains with 5000 step burn-in followed by 25000 samples, of which 5000 thinned samples were kept for subsequent analyses. We used these 5000 samples of each parameter to calculate the associated trait thermal curves, resulting in 5000 thermal fits of the trait data. Next, we use the mean of all 5000 posterior distributions as the thermal curve fitting the trait data.

After generating the 5000 posterior mean curves for each of trait, we derive the 5000 posterior curves for *R*_0_ then calculate the mean to get the temperature-dependent *R*_0_. For each posterior mean, we calculated the corresponding 95% highest posterior density (HPD) interval which is the smallest credible interval in which 95% of the distribution lies [32]. Our analysis was implemented in `R` [33]. More details on likelihoods and priors used can be found in Appendix S1.

### 2.3 Uncertainty in *R*_0_

The *R*_0_ formulas (Equation 1, 2, 3) depend on multiple temperature-sensitive traits, and so do their posterior densities. Hence, there are many sources of uncertainty in the mean posterior density that can be identified through uncertainty analysis. We calculate the uncertainty associated with *f, g*, and *V* by varying one while keeping the rest constant.

Next, we calculate the 95% credible interval around the mean posterior curve and measure the relative width of the interval, i.e. the difference between the upper and lower quantiles. We repeat this process for each individual component, *f, g*, and *V* then plot all the curves together against temperature. In this plot, we can identify which model component creates the most uncertainty in *R*_0_ by identifying the curve with the highest value at a given temperature.

### 2.4 Mapping suitability

To visualize and apply our understanding of the thermal suitability of BTV, we mapped both suitability and risk, at global scales. First, we present the geography of suitability across the globe by mapping the number of months of suitable temperatures for transmission - i.e. where *R*_0_ >0, using monthly average temperatures from the WorldClim dataset [34]. We note that here we use a scaled form of *R*_0_ by dividing the thermal performance of *R*_0_ by the maximum *R*_0_ value.

Second, we map livestock at risk of transmission, using the latest FAO Gridded Livestock of the World (GLW3) data for 2010, which details global distributions of sheep, goats, cows, and others, at a 5 minute scale [35]. To create a visually accessible risk map, suitability was scaled 0-1, and this was multiplied by 1+log(livestock). Thus we create a scaled risk map, balancing the season length and livestock density, to emphasize areas of coincidence, rather than simple suitability. In this case we used the GLW3 sheep distribution [36], as the primary host at risk. All map calculations and manipulations were run in `R` using packages `raster` [37, 38], `maptools` [39] and `Rgdal` [40], following methods described in [41].

## 3 Results

### 3.1 Temperature-dependence model components

Here we summarize the model components that depend on temperature and explain their role in the model.

#### Midge thermal traits

Figure 4 shows two columns, one for development time and one for survival probability. The development time column shows the time in days that eggs, larvae, and pupae need to develop to the next stage. The data are fitted using a quadratic function, under the assumption that juvenile midges at a given stage will need more time to develop at very low (<20°C) and very high (>35°C) temperatures. For eggs, the development time ranges from 60 to 70 days; for larvae, from 15 to 35 days; and for pupae between 40 and 80 days.

**Figure 4:**
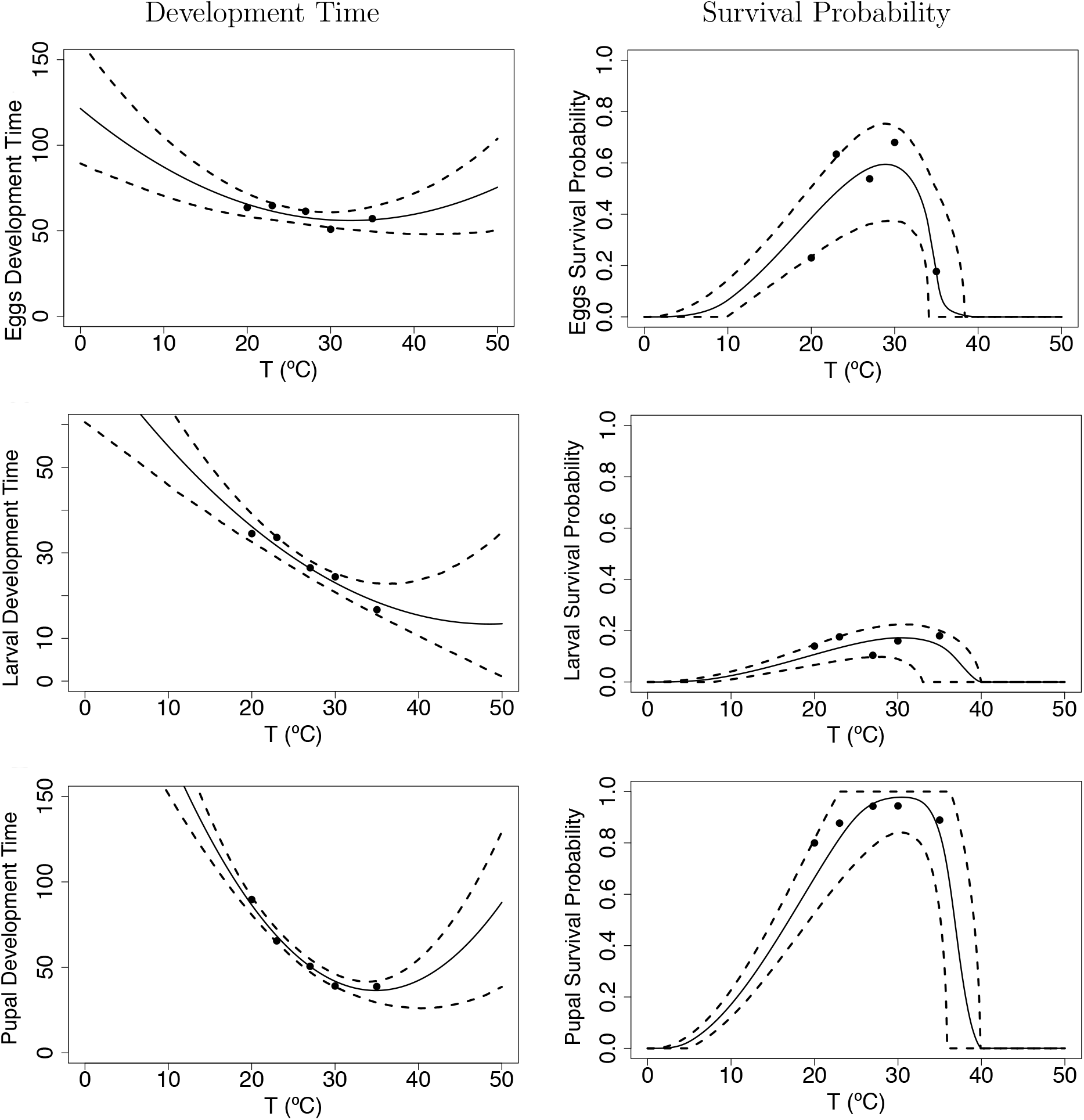
Figures in Column A show development time in days for midge juvenile stages, eggs *ρ_E_*, larvae *ρ_L_*, and pupae *ρ_P_*. Figures in Column B show survival probabilities for midge juvenile stages, eggs *p_E_*, larvae *p_L_*, and pupae *p_P_*. The solid line is the mean of the posterior distributions of the thermal response curves while the dashed lines represent the HPD intervals

On the other hand, the second column shows survival probabilities for eggs, larvae, and pupae. We fit the survival probabilities using a Brière curve (Figure 4 B). The survival probability is relatively high for eggs (0.2 < *p_E_* < 0.8), very low for larvae (*p_L_* < 0.2), but almost always 100% for the pupae stage (*p_P_* ~ 1). For fecundity *F*, we use data on the number of eggs laid per female per day. To fit *F* we use a Brière curve as shown in Figure 5 A. The fecundity reaches a maximum around 30°C and we do not have data for temperatures beyond that. The mortality rate, *μ*, is fit using a quadratic curve where we assume that the mortality is highest for temperature less than 10°C and higher than 30°C (Figure 5 B).

**Figure 5:**
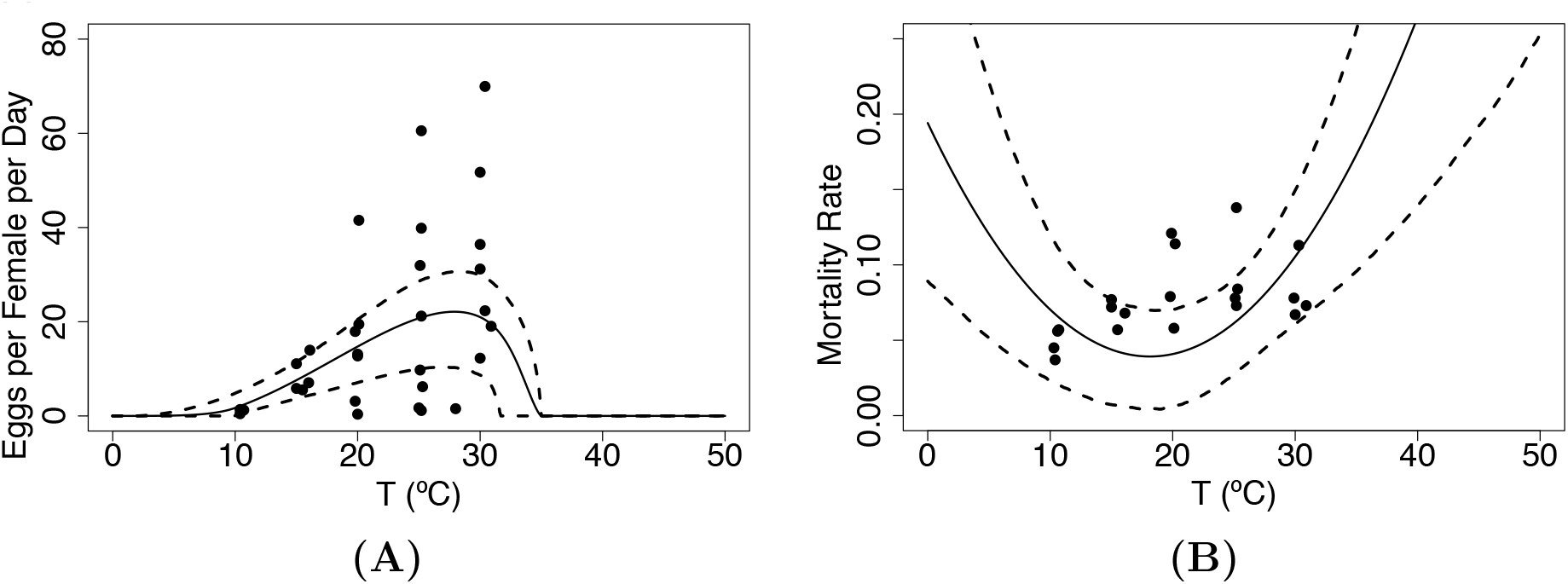
(A) Eggs per female per day *F* and (B) adult mortality rate *μ* traits as they vary with temperature. The solid line is the mean of the posterior distributions of the thermal response curves while the dashed lines represent the HPD intervals

Figure 6 shows the biting rate *a* and the transmission probability *b* both fit with a Brière curve. The biting rate minimal values lie around 10°C and increase to reach a maximum at 30°C. While the transmission probability *b* is minimal around 15°C and reaches a maximum at 30°C. We do not have data for the infection probability *c* so we assume that it is equal to 0.5. Lastly, Figure 7 shows the parasite development rate fit using a Brière curve, with minimal values around 15°C and maximal values around 32°C. Overall, these thermal traits all lack data values at extreme temperatures.

**Figure 6:**
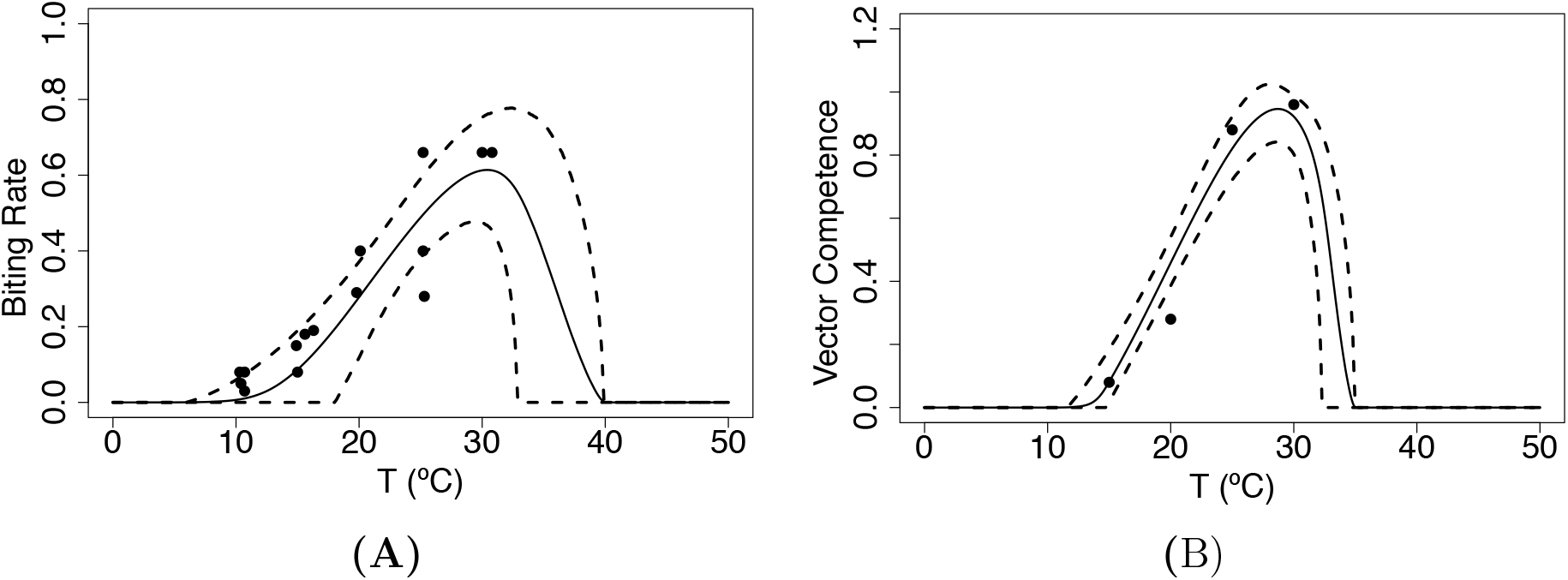
(A) Biting rate *a* and (B) probability that midges transmit infection when biting an uninfected host *b*. The solid line is the mean of the posterior distributions of the thermal response curves while the dashed lines represent the HPD intervals

**Figure 7:**
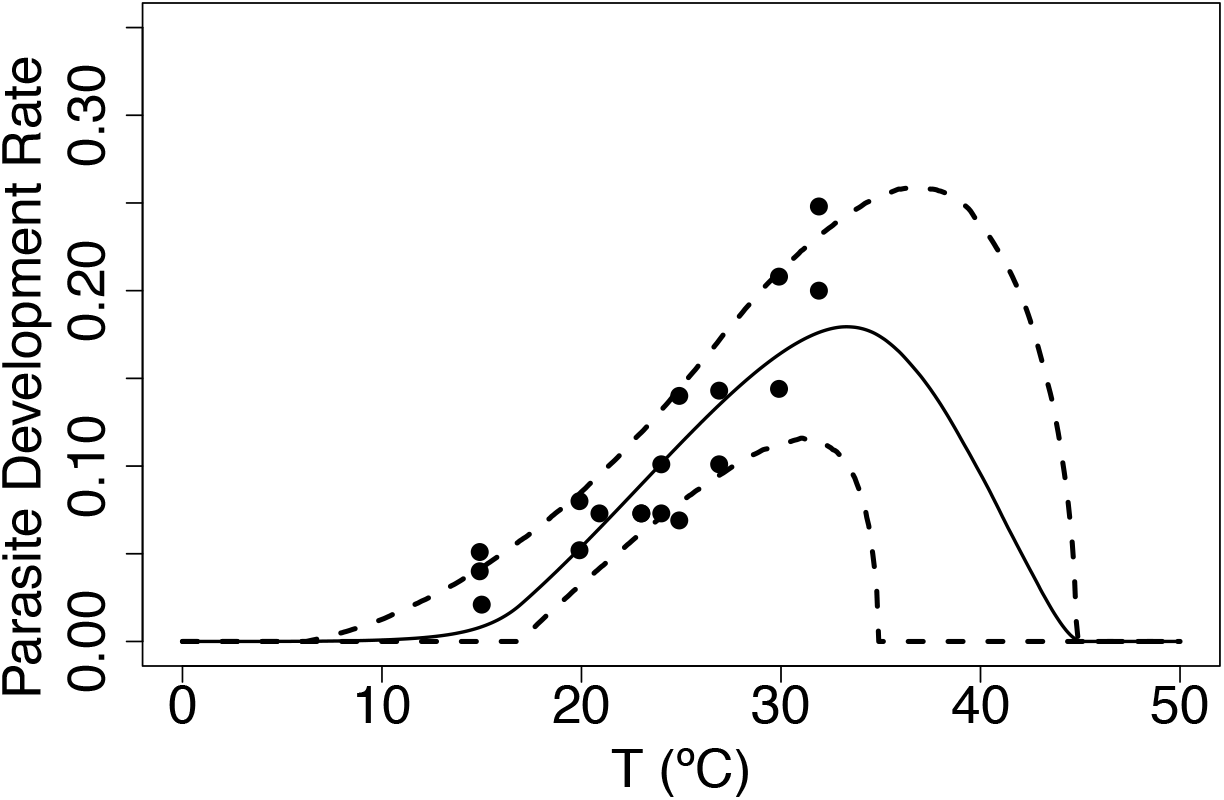
Parasite development rate (*ν*) is the inverse of extrinsic incubation period (*ν* = 1/*EIP*). The solid line is the mean of the posterior distributions of the thermal response curves while the dashed lines represent the HPD intervals

#### Midge density *V*

Recall the midge density formula given by

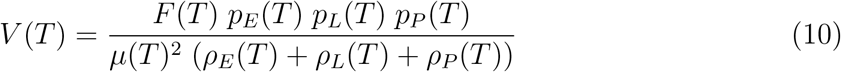

To estimate midge density *V*, we use survival probabilities *p_E_, p_L_, p_P_*, for egg, larvae and pupae; and development times *ρ_E_, ρ_L_, ρ_P_* corresponding to the egg, larvae, and pupae life stages; the fecundity measure represented by the number of eggs per female per day *F*; and adult mortality rate *μ*. Given all the traits contributing to the midge density formula we can evaluate *V*(*T*) to visualize how midge density varies with temperature. Figure 8 shows that midge density *V* is highest between 20 °C and 28 °C; it increases at temperatures higher that 10 °C and decreases when temperature exceeds 28 °C.

**Figure 8:**
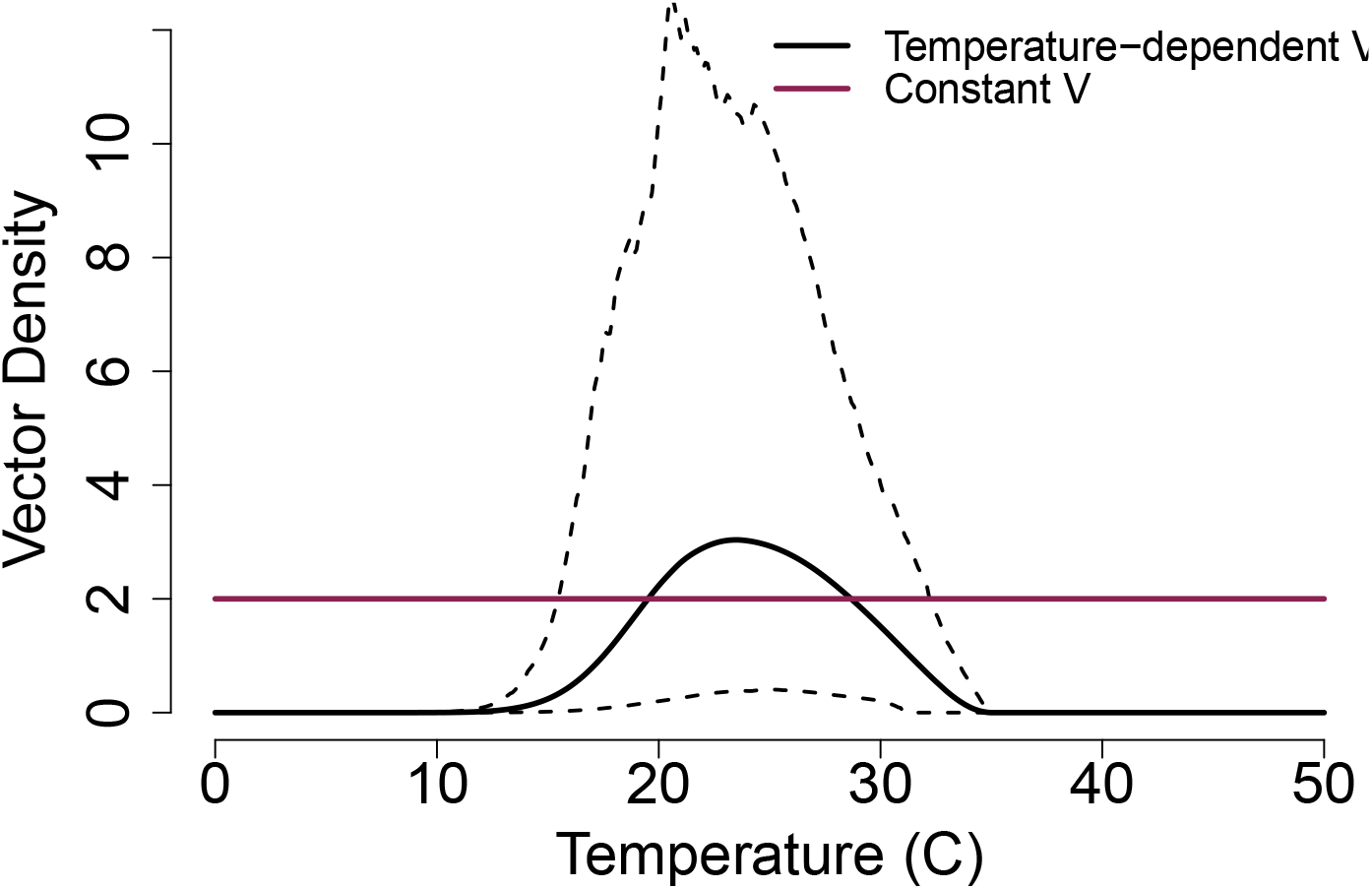
Modeled midge density as it varies with temperature. To obtain the temperature-dependent midge density, we evaluate Equation 10 at all temperature-dependent traits using the fitted curves. The solid black line shows the estimated density and the dashed lines show the corresponding HPD interval. A constant value *V* = 2 shown for comparison for subsequent modeling.

#### Transmission potential

The component *g*, is estimated by calculating the product of midge biting rate *a* and vector competence *bc*. For the transmission probability *c*, we assumed that there will be a 50% chance for midges to become infected after biting an infected host *c* = 0. 5. Figure 9 shows the transmission potential thermal curve and its linear dependence on *c*. Ideally, when data are available we suspect that *c* will also depend on temperature and would have a hump-shaped thermal curve as well.

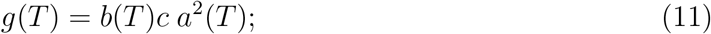

**Figure 9:**
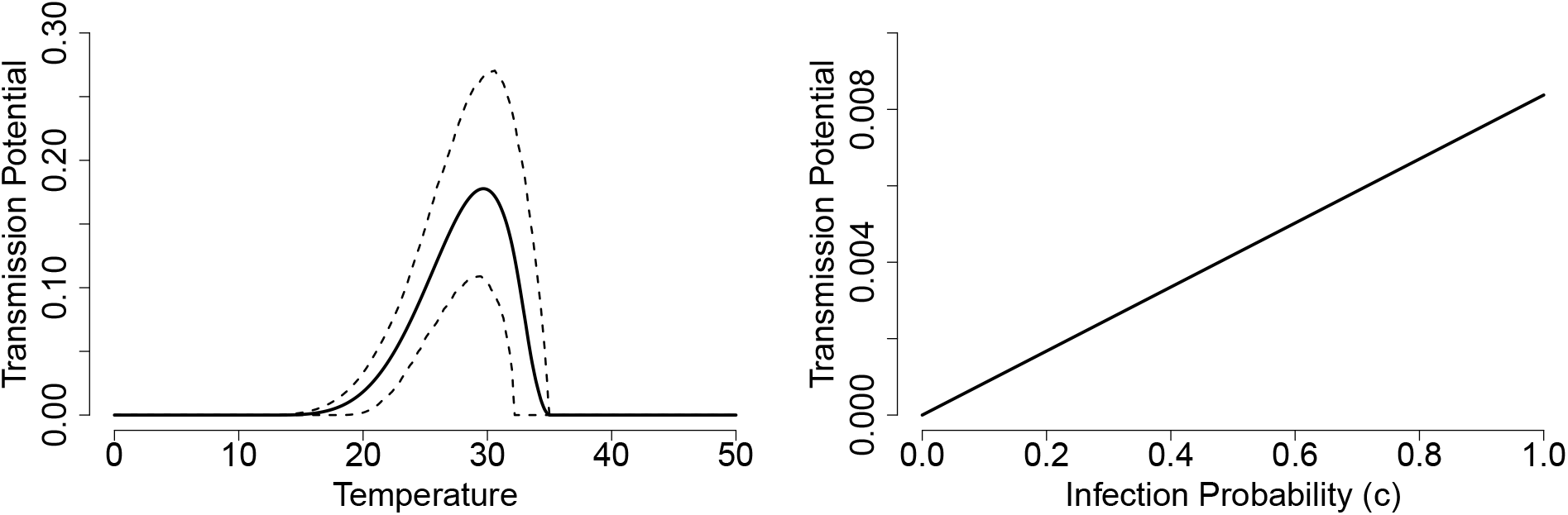
(Left) The transmission potential *g* as the biting rate *a* and transmission probability *b* vary with temperature while the infection probability is constant *c* = 0.5. The solid line shows the estimated curve and dashed lines are the HPD interval. (Right) We fix the biting rate *a* = mean(*a*(*T*) = 0.18 and transmission probability mean(*b*(*T*) = 0.23 and vary the infection probability between 0 and 1.

#### Functional form

Equation 12 shows all the different formulas used in the models we present in this paper. All three formulas use the mortality rate *μ* given in Figure 5 (B), and the parasite development rate *ν* fitted using a Brière curve in Figure 7. Recall, each of these formulas is the probability that a midge becomes infectious post infection. Figure 10 shows the variation of the functional form with temperature based on the two temperature-dependent traits *μ* and *ν*. Although there are differences between the magnitude of these curves, we can see that their peak occurs at the same temperature (25 °C), which is due to the traits’ thermal curves used in the formula. In addition, all of their HPD intervals overlap which means that there are no significant differences between them.

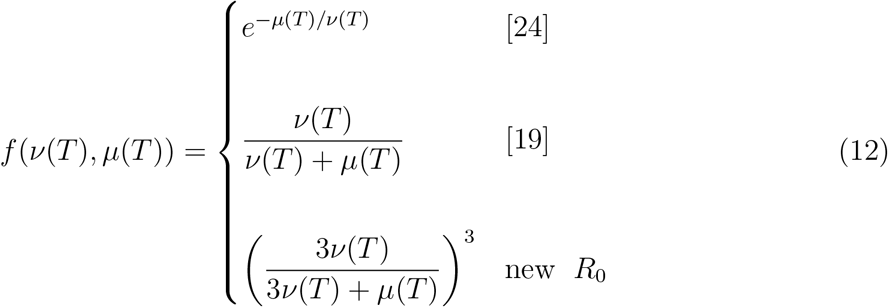

**Figure 10:**
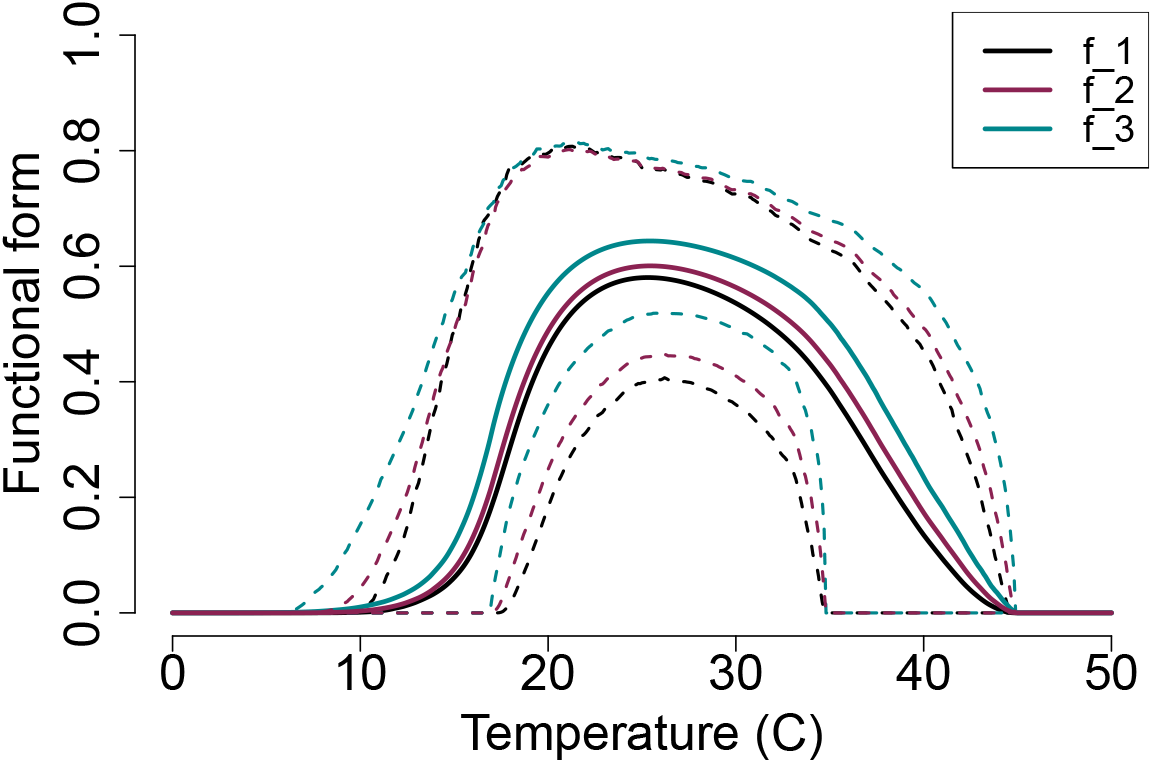
Functional forms used in *R_0_* versus temperature, the black line shows our model with the newly derived *R*_0_, the purple line shows the model presented in [19], and the blue line shows the model presented in [24]. Each solid line represents a different model and the dashed lines show the corresponding HPD intervals. We note that there is an overlap between all HPD intervals meaning that there are no statistically significant differences between these models.

### 3.2 Thermal response of *R*_0_

We use thermal traits to evaluate *R*_0_ given by Equation 4 with constant midge density *V* (Figure 11 Top) and with temperature-dependent midge density *V*(*T*) (Figure 11 Bottom). The three models are slightly different when constant midge density is used but become agree when temperature-dependent midge density is used. This is due to all the temperature-sensitive traits used to calculate *V*(*T*); however, this also leads to a higher uncertainty shown in the range of HPD interval in Figure 11 (Bottom). The lower thermal bound of the three posterior means are different by a magnitude of 1°C. However, the peak temperature and upper thermal limits are in agreement for all three models. With these results, we predict that *R*_0_ > 1 occurs at temperature greater than 15°C and less than 33°C, meaning that BTV is likely to cause an outbreak within this temperature range. We note that this prediction is based on assuming *c* = 0. 5 which may not be always true in reality.

**Figure 11:**
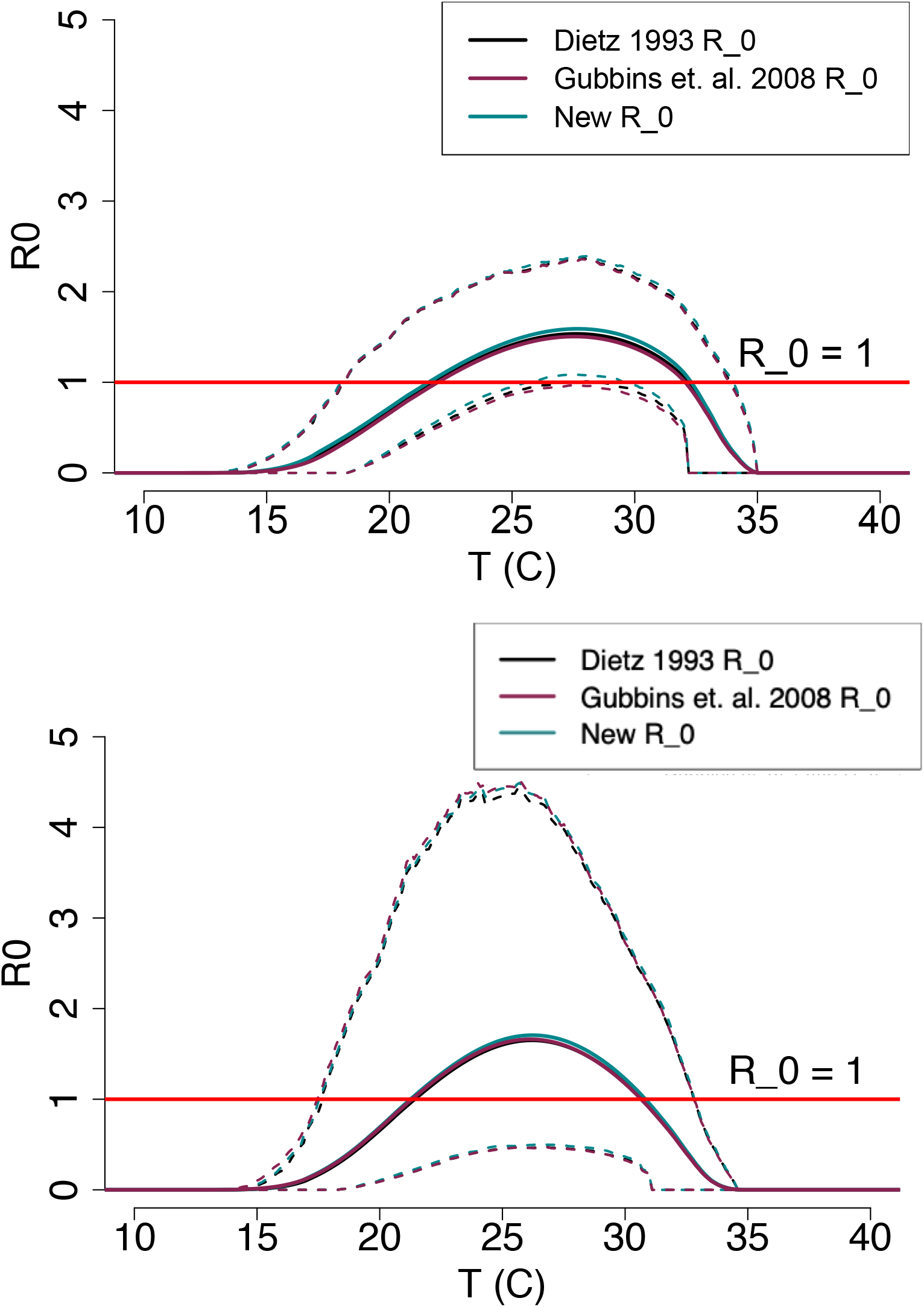
(Top) *R*_0_ with constant midge density *V* and (Bottom) *R*_0_ with temperature-dependent midge density *V*(*T*). The plots shows the magnitude of *R*_0_ changing as temperature increases. Each solid line represent the mean of the posterior distributions of *R*_0_ while the dashed lines are the HPD intervals.

### 3.3 Source of uncertainty in *R*_0_

In Figure 11 (Bottom), a high variation around *R*_0_ posterior density is shown in the large HPD interval. To determine the source of this uncertainty, we plot the calculated relative widths for each *R*_0_ component, see Figure 12. The results show that at a low temperature range (14°*C* < *T* < 18°*C*) uncertainty in *R*_0_ is mainly due to the uncertainty in the functional form *f*. At intermediate temperatures (18°*C* < *T* < 33°*C*), the uncertainty is caused by the midge density *V*(*T*). At very high temperatures (33°*C* < *T* < 35°*C*), the transmission potential *g* is the component producing the most variability in *R*_0_.

**Figure 12:**
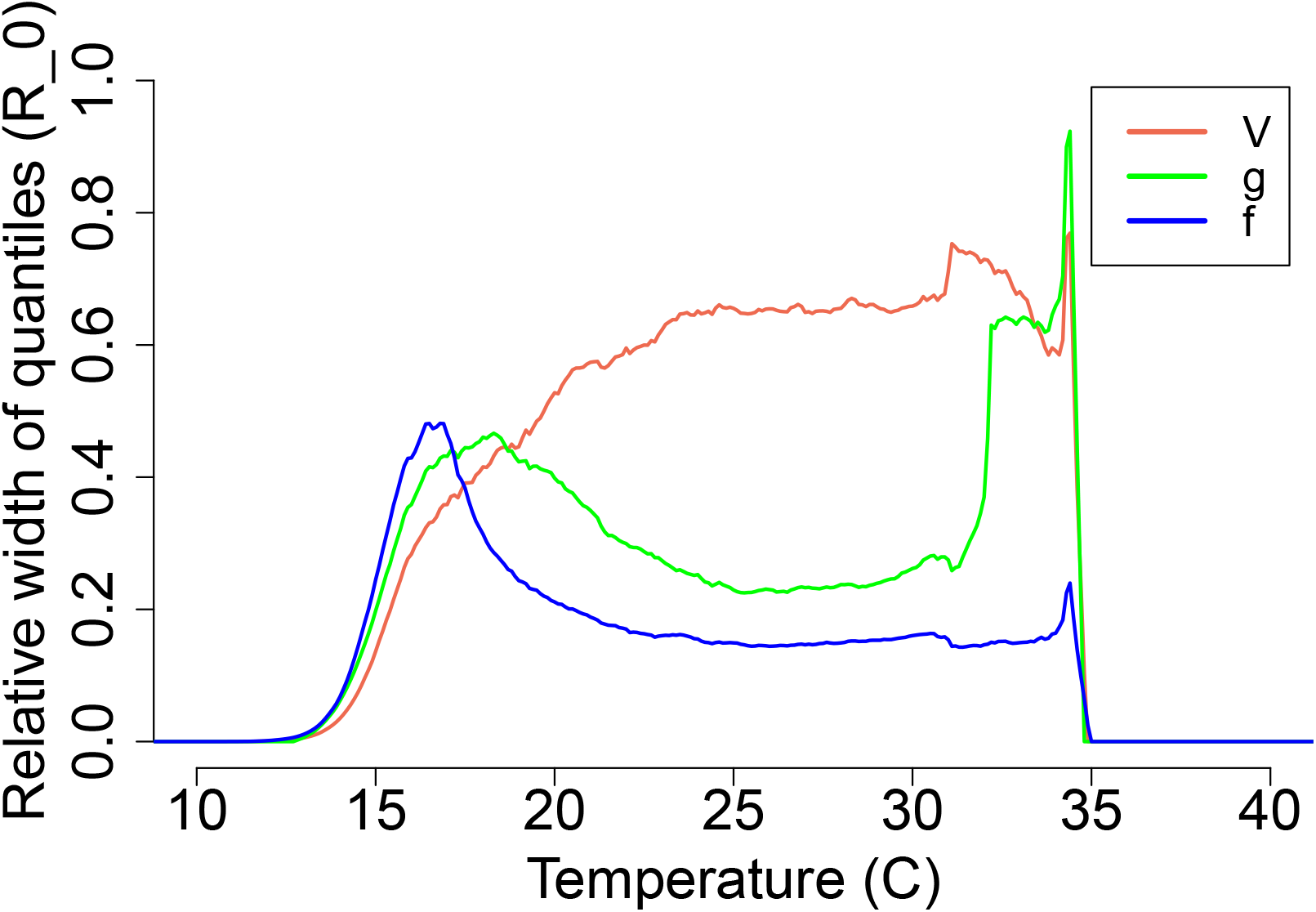
The source of uncertainty in *R*_0_(*T*) is measured by calculating the relative width of quantiles with each component varying with temperature while the remaining components are kept constants.

### 3.4 BTV risk maps

Figure 13 illustrates the number of months each area is at risk of BTV transmission with the assumption that *Culicoides sonorensis* and *Culicoides variipennis* are the main vectors. The results show that, under baseline long-term average current temperature conditions, much of central Africa, south Asia, central and the northern part of South America, and northern Australia are suitable for year-round bluetongue transmission. These areas are also the warmest parts of the world, and as we move away from them, the temperature is lower and the number of months of suitability is reduced.

**Figure 13:**
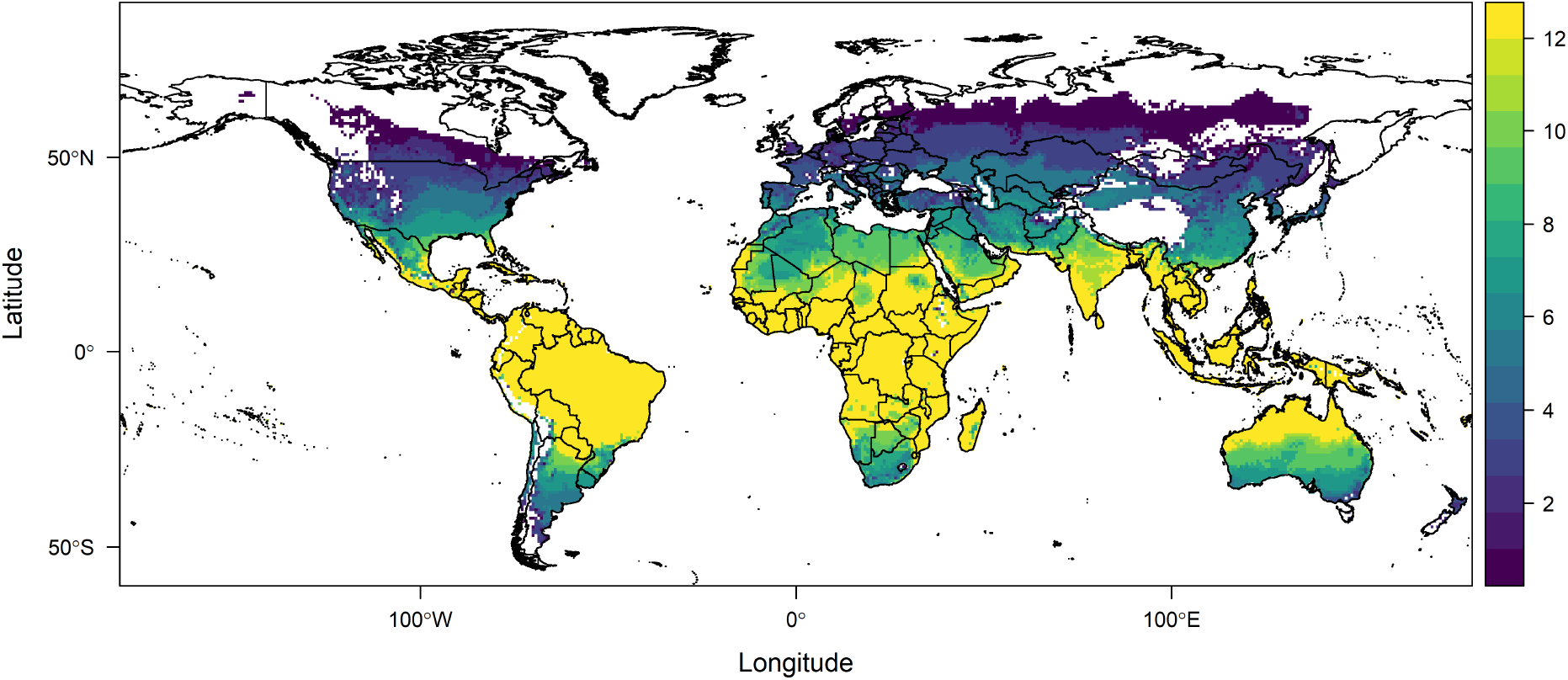
Map of the number of months (1-12) areas are at risk of bluetongue virus transmission according to our temperature-dependent *R*_0_. This map based on the current mean monthly temperatures and is restricted to bluetongue disease caused by the two midge species *Culicoides sonorensis* and *Culicoides variipennis*.

Next, we used the global distribution of sheep to determine areas where sheep are at risk of acquiring BTV. The choice of sheep was mainly due to ready data availability, and also because sheep are the BTV host with highest mortality and morbidity rates, and therefore of great interest and relevance. The map shows that areas where sheep are at the highest risk (scale > 3) are located around the equator. The next highest risk regions (1 < scale < 3) are areas with of high livestock industry, such as central and south America and Europe.

**Figure 14:**
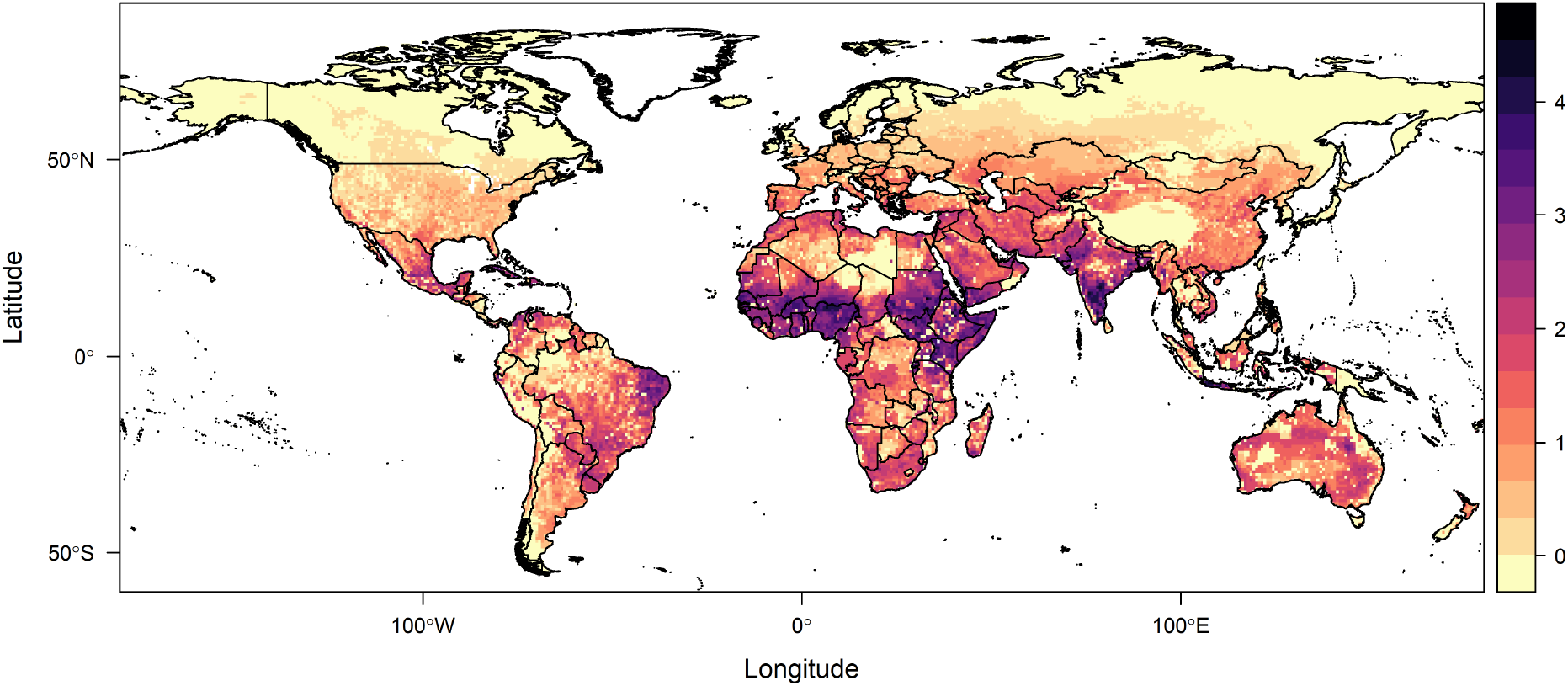
Scaled transmission risk suitability of bluetongue virus for sheep, as the primary host at risk, worldwide. the scale ranges from a low risk, 0, to a high risk, 5.

## 4 Discussion

Here we are interested in the effect of temperature on BTV spread. As highlighted in a 2018 systematic review [42], BTV has been studied using many different modeling approaches. The systematic review summarized BTV models used post-1998 [42], most of which relied more on strong modeling assumptions than data. The model results were used to inform animal health decision-making by identifying at-risk areas and the risk of spread in case of introduction [43]. Although a few models have used the *R*_0_ approach to study BTV, our model differs in that it incorporates temperature across a wide range, leading to estimating an *R*_0_ that is also temperature dependent. Linking *R*_0_ to temperature can help identify BTV outbreak risk based on temperature at particular locations. This can inform management policies and control strategies, within current and changing climate conditions.

We use a Ross-Macdonald type modeling approach to describe BTV transmission [22, 23]. We also adopt two previously used BTV models, [24] and [19], to compare their resulting *R*_0_*s* to ours. This mechanistic approach allowed us to derive the basic reproduction ratio’s posterior distribution as a function of temperature. We were able to both determine the suitable temperature for possible BTV outbreaks when *R*_0_ > 1 and the temperatures at which BTV outbreaks are likely to die out when *R*_0_ < 1. We note that the thermal response of *R*_0_ here is dependent on our model assumptions, for example setting the infection probability to be *c* = 0.5 and choosing the host density *H* and recovery rate *d* such that *Hd* =1.

Based on the available traits’ laboratory data we used in our model, we predict that BTV outbreaks, at least from the examined midge species, can occur within the temperature range of 15°C and 33°C, with outbreak peak at about 26°C. This result was obtained regardless of which *R*_0_ formula used, i.e., all three different models lead to the same predictions. Similarly, investigating the lower thermal limit, peak, and upper thermal limit of *R*_0_ led to no difference between peaks and upper thermal limits for all three forms, and a difference between lower thermal limits of only 1 °C. This indicates that the uncertainty of temperature effects outweigh the effects of differences in modeling assumptions for these models.

Previous studies have investigated temperatures suitable for other vector-borne diseases. For example, a study on three mosquito-borne diseases, Zika, dengue, and chickungunya transmitted by *Aedes aegypti* and *Aedes albopictus* showed that the transmission is likely to occur between 18-34°C with peak transmission between 26-29°C [44]. Moreover, the temperatures suitable for the transmission of the plant-borne disease, citrus greening, between 16°C and 33°C with peak transmission at 25°C [45]. Together with our findings, this shows that there are similarities between ectotherm vectors in the way they respond to temperature. For example, their traits follow humped shaped thermal performance curves. But there are differences in the temperatures ranges they tolerate, and the temperatures at which their performance is maximal.

Incorporating different temperature-dependent traits into the model led to uncertainty in the *R*_0_ posterior density. Our uncertainty analysis allowed us to determine the traits responsible for causing uncertainty in *R*_0_ at each part of the temperature range. At a low temperature range (14°*C* < *T* < 18°*C*) more data are needed for the parasite development rate, *ν*, and mortality rate, *μ*, to reduce this uncertainty in the functional form *f*. At moderate temperatures (18°*C* < *T* < 33°*C*) the uncertainty in *R*_0_ is caused by *V*, meaning that more data are needed in traits contributing to estimating the midge density. The biting midge density *V*(*T*) appears to be the main source of uncertainty across a wide temperature range. At very high temperatures (33°*C* < *T* < 35°*C*) we need more data on vector competence *bc* and biting rate *a*. Reducing the uncertainty in *R*_0_ will refine our predictions and our control and prevention suggestions.

We were interested in using our derived *R*_0_ thermal response to determine areas at risk of BTV. A straightforward risk map can be a useful planning tool, both to understand the scale of current risk, and to anticipate suitable regions where establishment of BTV could be successful were it to be introduced, with competent vectors. To omit the effect of our parameters lacking data, namely, host density *H*, host recovery rate *d*, and infection probability *c*, we used a scaled form of *R*_0_ for spatial analyses and visualization. We divide the thermal response of *R*_0_ by the maximum value of *R*_0_ which leads to a temperature varying *R*_0_ curve with values between 0 and 1. We created global risk maps showing the number of months per year each location worldwide is suitable for BTV disease transmission given the presence of two midge species, namely, *Culicoides sonorensis* and *Culicoides variipennis*. The results show that warmer areas are at risk year-round, while cooler areas are at risk for fewer months. However, these results are based on long-term, baseline current temperatures, and with climate change, and the continuous rising of global temperatures, the area at risk of BTV may expand and shift to include places with previously lower risk [46].

## Acknowledgement

L.R.J. was supported by the National Science Foundation Career grant (NSF DMSDEB #1750113) and the National Institutes of Health, NIH-USDA, Ecology of Infectious Diseases (Award Number 1518681).

## Authors’ Contributions

F.E.M. and L.R.J. designed the study. F.E.M., H.S., and Z.T. collected the data from the literature and performed the Bayesian analyses. F.E.M and L.R.J. performed the mathematical analyses and S.J.R. performed the spatial mapping. F.E.M. wrote the paper. All authors contributed to revising and editing the paper and gave approval for publication..

## S1 Appendix

### S1.1 Transmission model for BTV

We use an SIR-SEI type of compartmental model to describe vector-host interactions in transmitting BTV (see Figure 2). The host population (*H*) is divided into susceptible (*S*), infected (*I*), and recovered (or immune) (*R*) classes, while the vector population (*V*) is divided into susceptible (*S_V_*) and infected (*I_V_*) classes as well as three Exposed (*E_V_*) classes. Here we use three exposed classes in the vector population to incorporate a more realistic length of the extrinsic incubation period. Using three compartments with the exit rate from each compartment being 3 *ν*, lead to a Gamma distribution for overall midge progression to the infectious class with a mean rate of *ν*. Increasing the number of compartments used from 3 to a larger number leads to a Gamma distribution with lower variance around the mean [47]. This approach is an alternative to using fixed time delays, which are not suitable when using temperature-dependent parameters. Both host (*H*) and vector (*V*) populations are assumed (and are by definition of the model) constant.

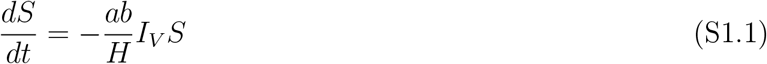

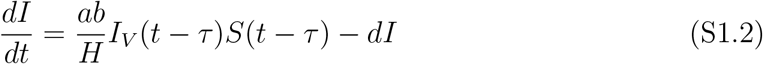

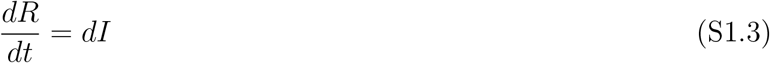

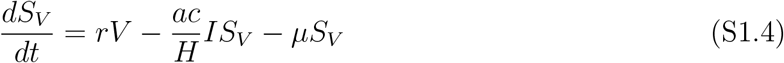

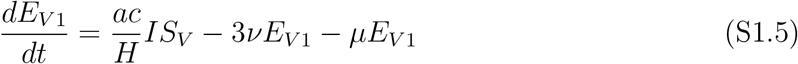

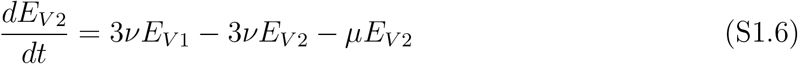

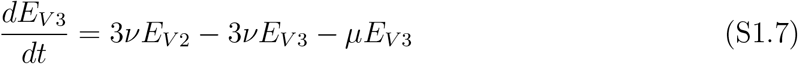

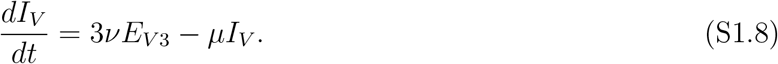

where

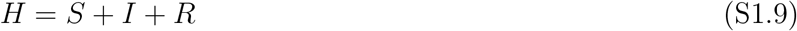

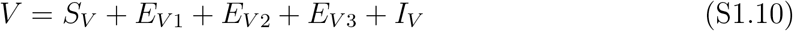

The model’s parameters are presented in the Table S1.1 below. Note that the parameters *τ* is the time that a susceptible (*S*) takes to become infected after receiving a bite from infected vector (*I_V_*). The parameter *ν* is the inverse of the Extrinsic Incubation Period of the pathogen (EIP), i.e., 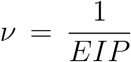. We define the vector population size to be 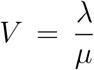, where *λ* is the total birth rate of adult midges in the whole population (adults/day), and *μ* is per-capita adult mortality rate (1/day). This is based on Parham & Michael [48], who derive the expression phenomenologically by treating *V* as a random variable. Thus, *λ* is equivalent to *rV* in the model above, given that *r* = *μ* at disease free equilibrium, and is given by

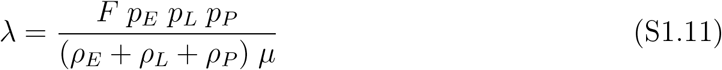

where *F* is the number of eggs produced by all females in the population per day, *p_E,L,P_* are the survival probabilities in the Eggs, Larvae, and Pupae stages, and *ρ_E_, ρ_L_, ρ_P_* are the development time in each stage. Then, the abundance of the vector becomes,

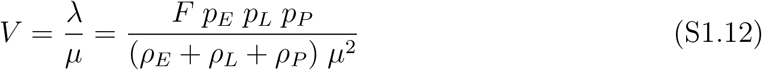

**Table S1.1:** Parameters used in the mathematical model, their description, and units.

| Parameter | Definition | Units |
| --- | --- | --- |
| $d$ | Recovery rate of infected hosts | 1/day |
| $\tau$ | Host's exposure period | day |
| $r$ | Vector population's birth rate | 1/day |
| $a$ | Vector biting rate | bites / day |
| $b$ | Probability that a midge is infected | dimensionless |
| $c$ | Probability that a midge is infectious | dimensionless |
| $\mu$ | Mortality rate of adult vectors | 1/day |
| $\nu$ | Parasite's development rate | 1/day |

### S1.2 Host recovery rate *d* sensitivity analysis

Although all ruminants are susceptible to BTV disease, each responds to the infection differently, with sheep being the most susceptible and showing extreme morbidity and mortality. In addition, BTV host recovery depends on the intensity of the infection as well as the time of disease detection, which results in recovery rate variability among hosts. To account for this, we perform a sensitivity analysis on the host recovery rate *d* by looking at the derivative of *R*_0_ with respect to *d* as follows:

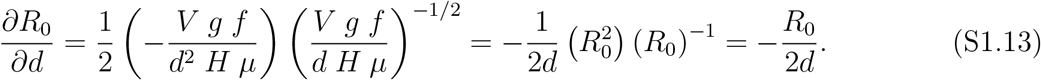

Since 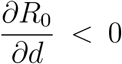 always, the basic reproductive ratio *R*_0_ increases as the recovery rate *d* decreases. Figure S1.1 shows different *R*_0_ densities corresponding to different host recovery rate values. Higher lengths of infection 1/*d*, i.e. lower recovery rates *d*, are associated with higher *R*_0_ densities, meaning that hosts with low recovery rate such as sheep are more challenging to manage as the chance of outbreak for them is more likely.

**Figure S1.1:**
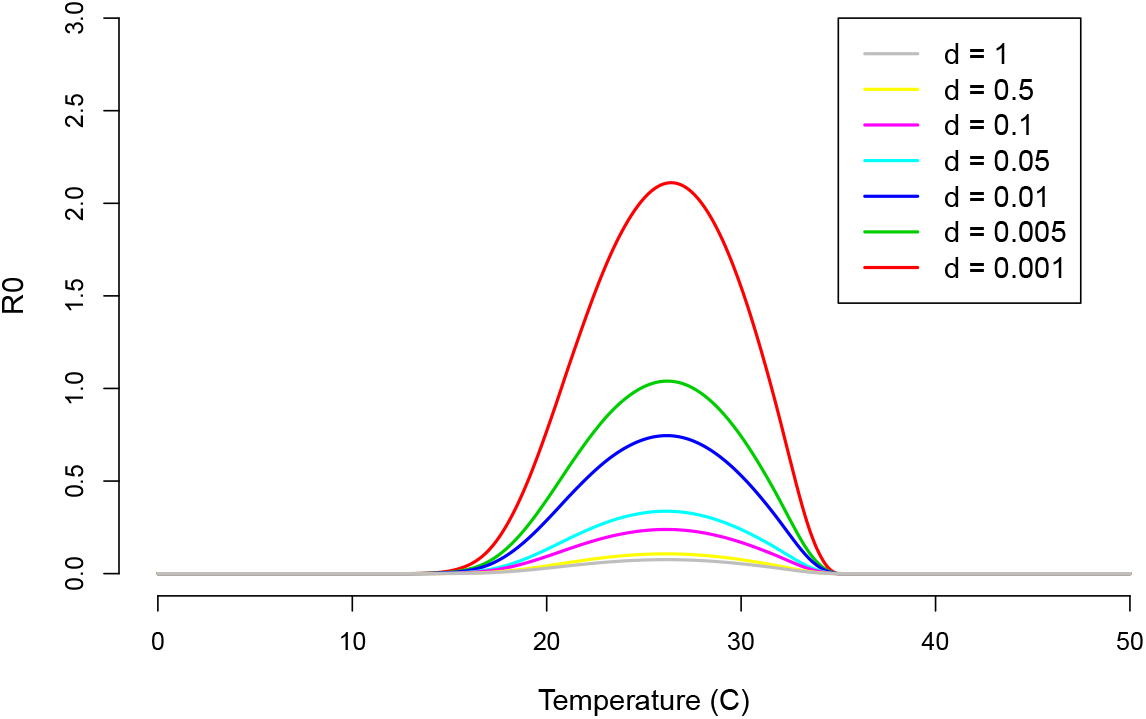
Host recovery rate *d* values correspond to different *R*_0_ posterior densities. As *d* decreases, *R*_0_ density increases meaning that a lower recovery rate correspond to a higher outbreak risk.

### S1.3 Uncertainty analysis

We investigate the uncertainty caused in each of *R*_0_ components, the midge density *V*, the functional form *f*, and the transmission potential *g* by examining the source of uncertainty within each component. For the midge density, the uncertainty is mainly caused by the adult midge mortality rate *μ* within a wide temperature range, from 10°C to 32°C. At higher temperatures (¿32°C) the uncertainty is caused by the fecundity *F*.

In the functional form case, the uncertainty is caused by the adult mortality rate *μ* for temperatures between 18°C and 32°C, this range overlaps with that of the midge density. At lower lower (10-18°C) and higher (32-45°C) temperature ranges, the uncertainty is caused by the pathogen development rate *ν*. In the transmission potential the overall uncertainty is caused by the biting rate *a*.

**Figure S1.2:**
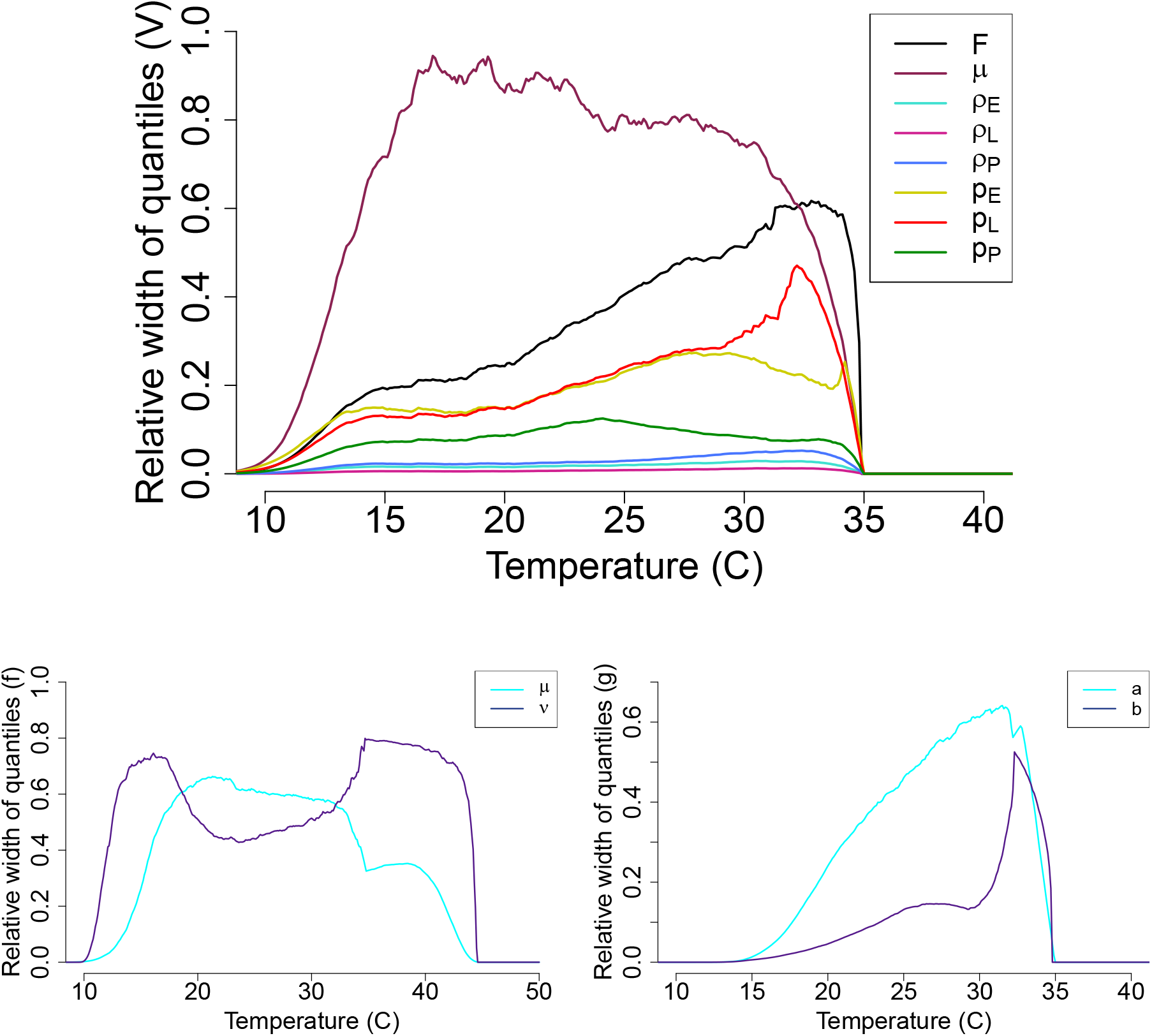
The source of uncertainty in the midge density *V* (Top), the functional form *f* (Bottom-left), and the transmission potential *g* (Bottom-right) is measured by calculating the relative width of quantiles with each parameter varying with temperature while the remaining parameters are kept constants.

### S1.4 Bayesian fitting of traits thermal curves

To fit each trait, we chose a unimodal functional form as the mean function. We use normal distributions for most of the data while binomial distributions are used when fitting probability distributions. We used uninformative priors appropriate for the biological description of the data, taking into account the positivity of their values as well as their range. The values in the priors are decided as we go until the appropriate fitting curve is obtained.

#### Midges biting rate *a*

The biting rate of adult midges is one of many factors that influence Bluetongue transmission [49]. In order to calculate the biting rate, the time required for female *Culicoides sonorensis* to lay eggs after a blood meal, also known as a gonotrophic period, is required. Biting rate (a) can be approximated by taking the inverse of the gonotrophic cycle duration. Similar to other traits, the biting rate is sensitive to environmental factors, especially, temperature [49] (see Figure S1.3 for thermal fit).

#### Vector competence *bc*

Vector competence for adult midges is a measure of their ability to transmit the disease. It is genetically determined and heavily influenced by environmental factors such as temperature and humidity [50]. Vector competence (bc) is the product of the probability of a vector getting infected after a blood meal containing a pathogen (c) and the probability of a vector transmiting infection (b). While we were able to find data for *b* concerning *Culicoides sonorensis* [51], we were unable to find data for *c*. We assume *c* = 0.5 for all calculations used in this analysis. We did fit a Bayesian model to the parameter *b* (Figure S1.4).

#### Juvenile survival probability *p_E_, p_L_, p_P_*

Vaughan et. al. studied the sub-adult life cycle of *Culicoides variipennis* at temperatures of 20 °C, 25 °C, and 28 °C [52]. We define the probability of an egg hatching by using the mean percentage of laid eggs that hatched at each given temperature. We now define the probability of successful larval pupation by collecting the percentage of larva that ended up pupating at each given temperature. We finally define the probability of pupae emerging to become adults, *p_P_*, as the mean percentage of pupae that survive to the adult stage at each given temperature (Figures S1.5, S1.6, S1.7).

#### Juvenile development time *ρ_E_, ρ_L_, ρ_P_*

Egg Development Time is defined as the time in days required for eggs to hatch in a given temperature. *Culicoides variiennis* were studied in a laboratory setting [52]. Larva Development Time is defined as the time in days required for the larva to mature into a pupa in a given temperature. Pupa Development Time is defined as the time in days required for a pupa to mature into adult midges in a given temperature (Figures S1.8, S1.9, S1.10).

**Figure S1.3:**
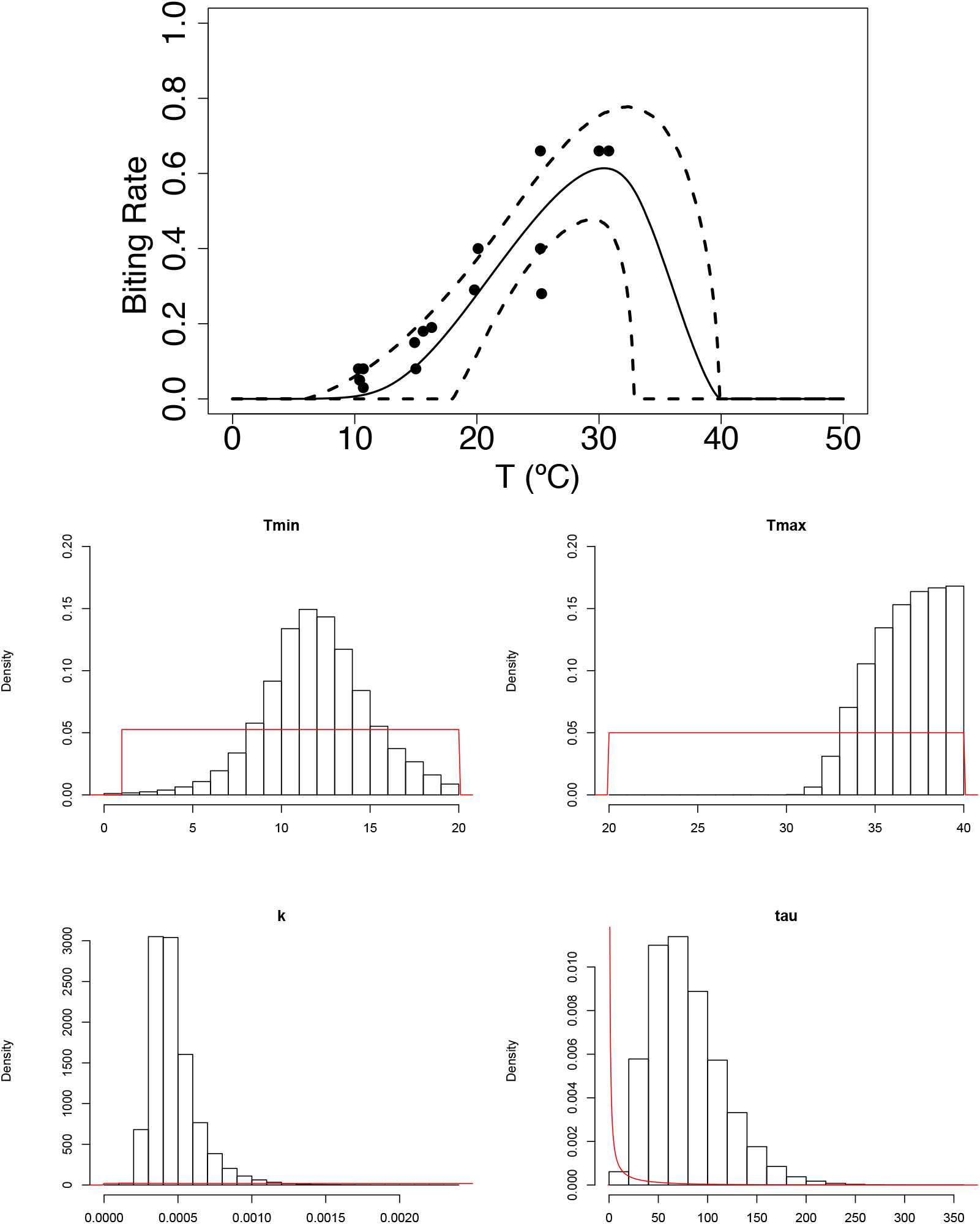
(Top) The mean trajectory in solid line and HPD interval in dashed black for the biting rate *a*. (Bottom) Histograms of the posterior distribution for each parameter of the Brière fit for the biting rate *a*. The prior distribution for each parameter is plotted in red. The Brière fit is determined by the equation 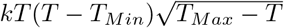 using a normal distribution with precision *τ*.

#### Fecundity *F*

The rate at which female midges lay eggs is closely related to the spread of Bluetongue. This rate is typically measured as eggs per female per day. For this study we also utilized fecundity data that was taken over two oviposition cycles and transformed the data (originally eggs per female) by dividing by the median oviposition time [49].

**Figure S1.4:**
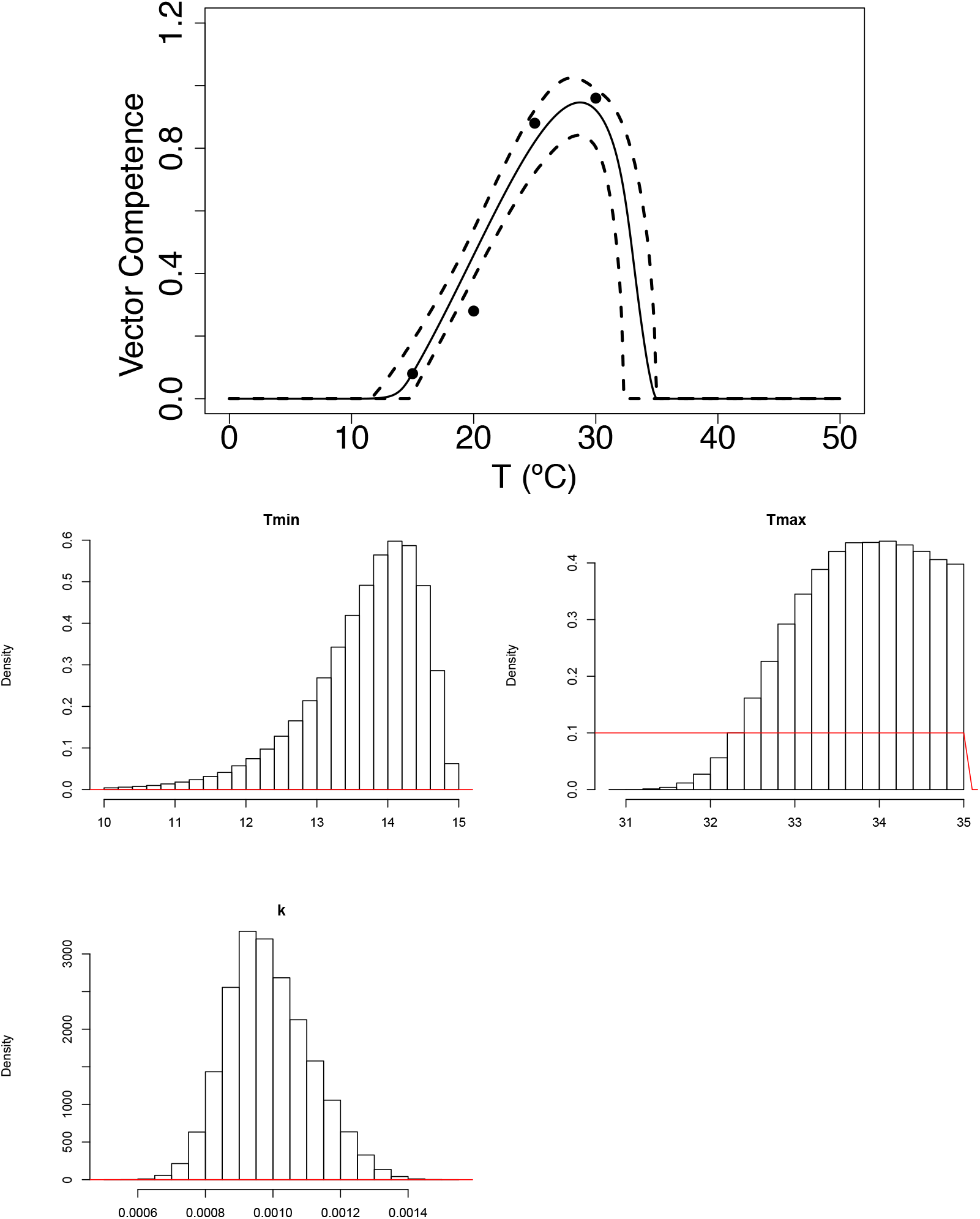
(Top) The mean trajectory in solid line and HPD interval in dashed black for the probability of a vector transmitting the virus when biting *b*. (Bottom) Histograms of the posterior distribution for each parameter of the Brière fit for the probability *b*. The prior distribution for each parameter is plotted in red. The Brière fit is determined by the equation 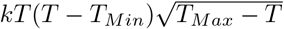 using a binomial distribution.

**Figure S1.5:**
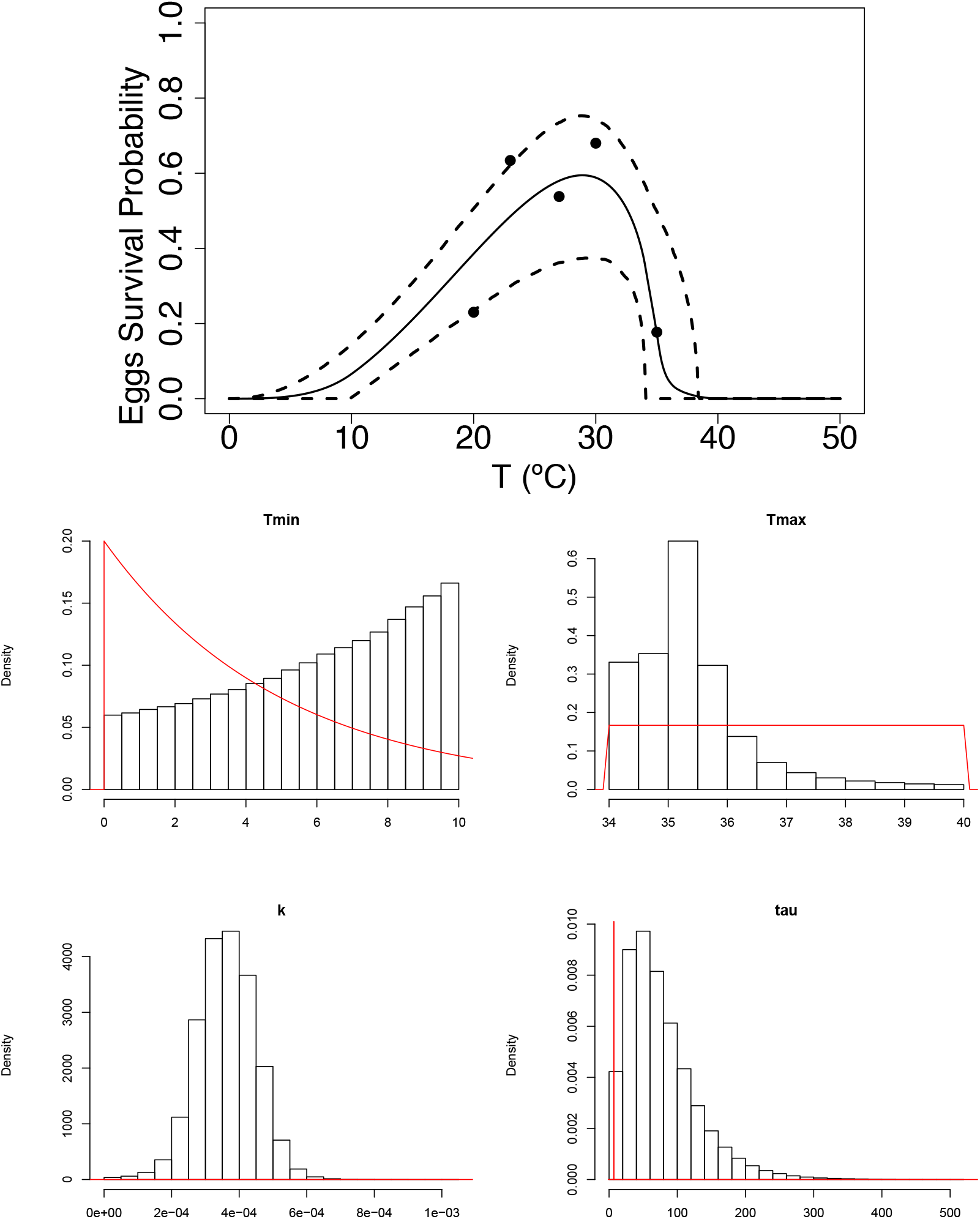
(Top) The mean trajectory in solid line and HPD interval in dashed black for the egg survival probability *p_E_*. (Bottom) Histograms of the posterior distribution for each parameter of the Brière fit for the probability *p_E_*. The prior distribution for each parameter is plotted in red. The Brière fit is determined by the equation 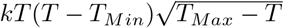 using a normal distribution with precision *τ*.

#### Pathogen development rate *ν*

Parasite development has been shown to increase with temperature in studies that support the hypothesis that global warming has been cause for latitudinal shifts which in turn increase the reach of vectors that transmit diseases like bluetongue [53]. In order to investigate this trait’s relationship with temperature, we made use of data on Extrinsic Incubation Period (EIP) to create a new parameter: Parasite Development Rate (*ν*) (*ν* = 1/EIP). EIP is the time between a vector getting infected with a pathogen to the time that the vector itself is able to transmit the pathogen.

**Figure S1.6:**
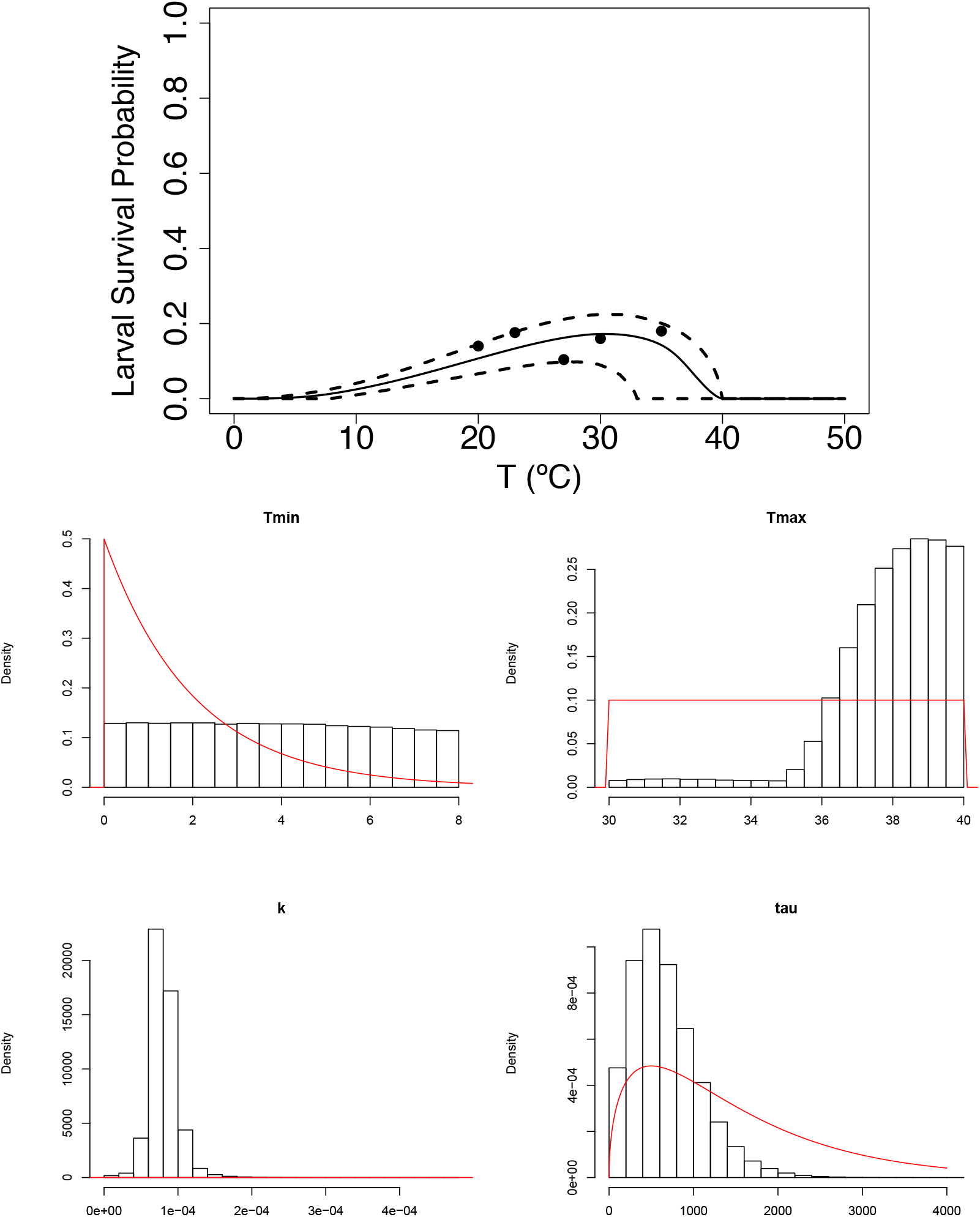
(Top) The mean trajectory in solid line and HPD interval in dashed black for the larval survival probability *p_P_*. (Bottom) Histograms of the posterior distribution for each parameter of the Brière fit for the probability *p_P_*. The prior distribution for each parameter is plotted in red. The Brière fit is determined by the equation 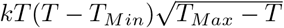 using a normal distribution with precision *τ*.

**Figure S1.7:**
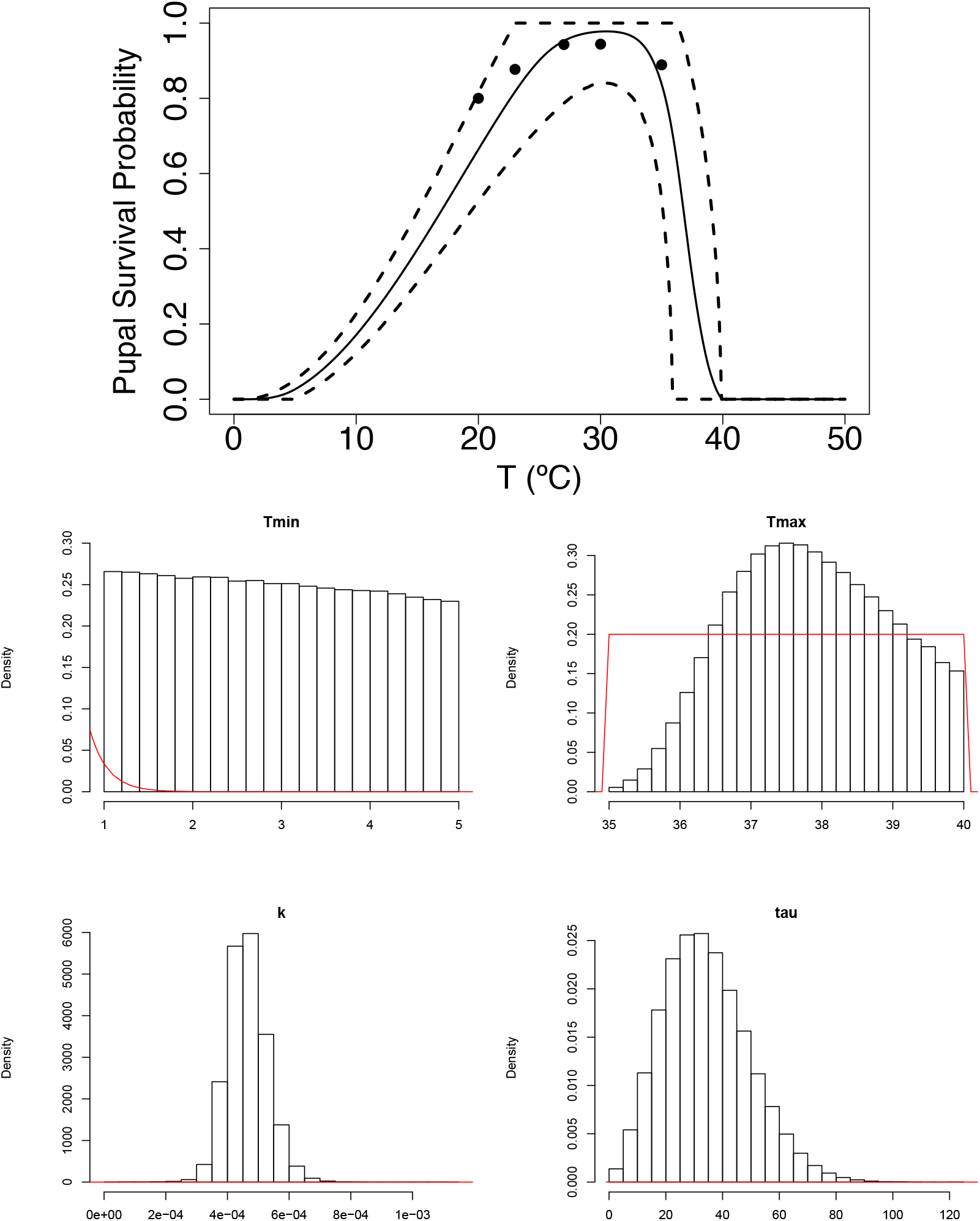
(Top) The mean trajectory in solid line and HPD interval in dashed black for the pupal survival probability *p_P_*. (Bottom) Histograms of the posterior distribution for each parameter of the Brière fit for the probability *p_P_*. The prior distribution for each parameter is plotted in red. The Brière fit is determined by the equation 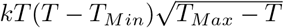 using a normal distribution with precision *τ*.

**Figure S1.8:**
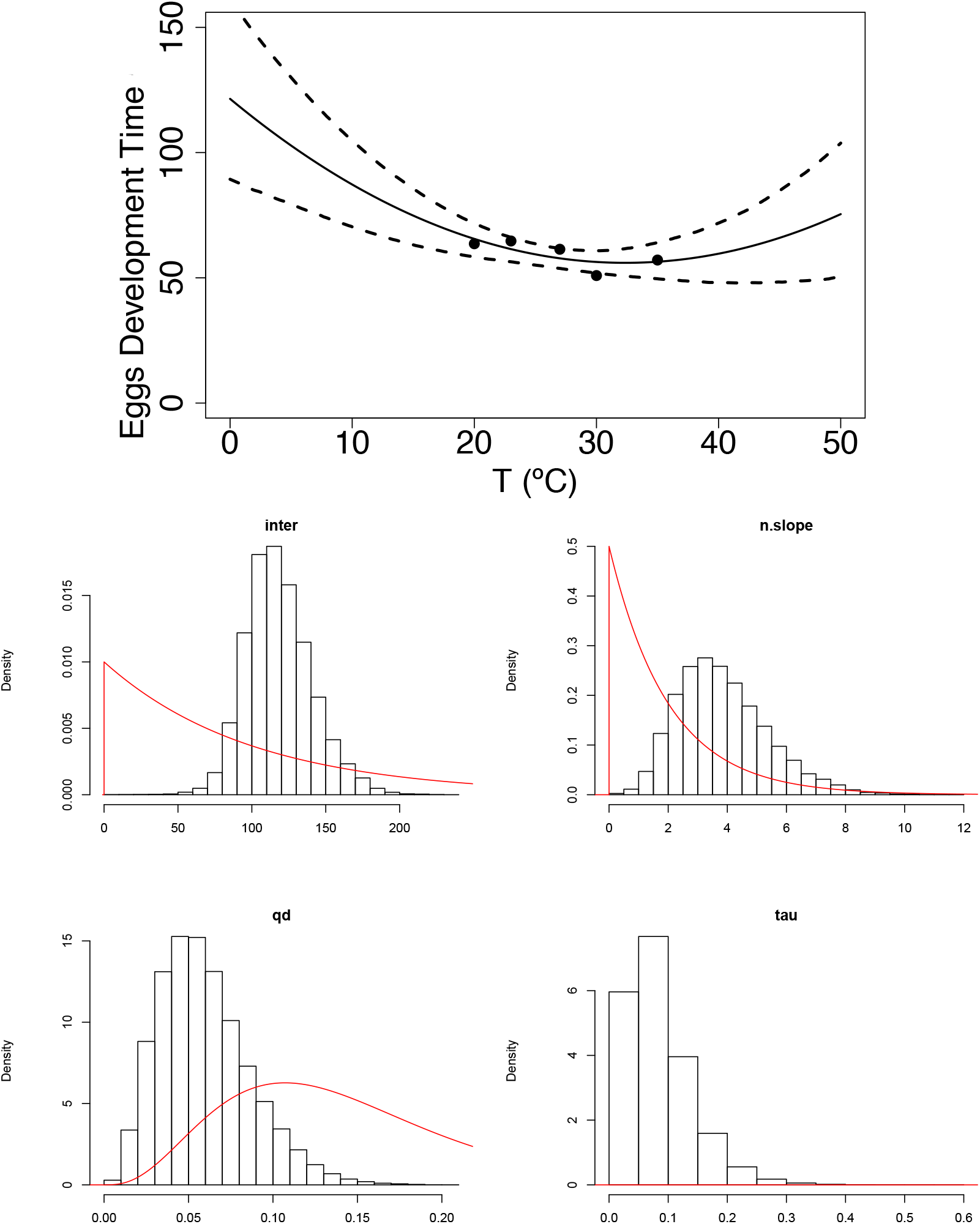
(Top) The mean trajectory in solid line and HPD interval in dashed black for egg development time *ρ_E_*. (Bottom) Histograms of the posterior distribution for each parameter of the quadratic fit for egg development time *ρ_E_*. The prior distribution for each parameter is plotted in red. The quadratic fit is determined by the equation *inter – n.slope T* + *qd T*^2^ using a normal distribution with precision *τ*.

#### Adult mortality rate *μ*

The rate at which midges die over a span of time is known as the mortality rate *μ*. We define the mortality rate of midges as 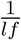, where *lf* represents the lifespan of midges in days, or the probability of survival for the midges. We define mortality rate in the case where *lf* is the lifespan of midges in days. Mortality rate is also sensitive to environmental factors, especially temperature [49].

**Figure S1.9:**
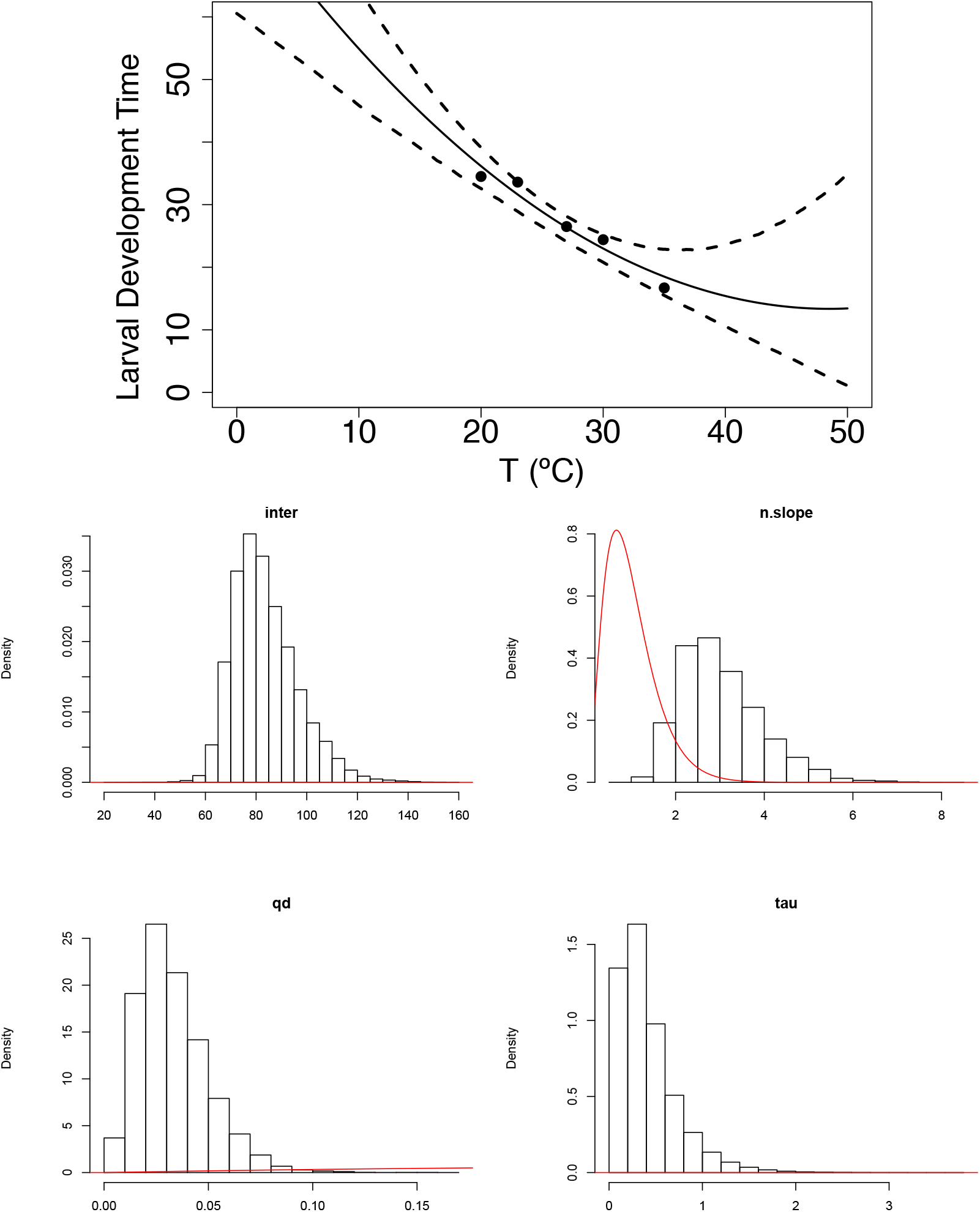
(Top) The mean trajectory in solid line and HPD interval in dashed black for larval development time *ρ_L_*. (Bottom) Histograms of the posterior distribution for each parameter of the quadratic fit for larval development time *ρ_L_*. The prior distribution for each parameter is plotted in red. The quadratic fit is determined by the equation *inter – n.slope T* + *qd T*^2^ using a normal distribution with precision *τ*.

**Figure S1.10:**
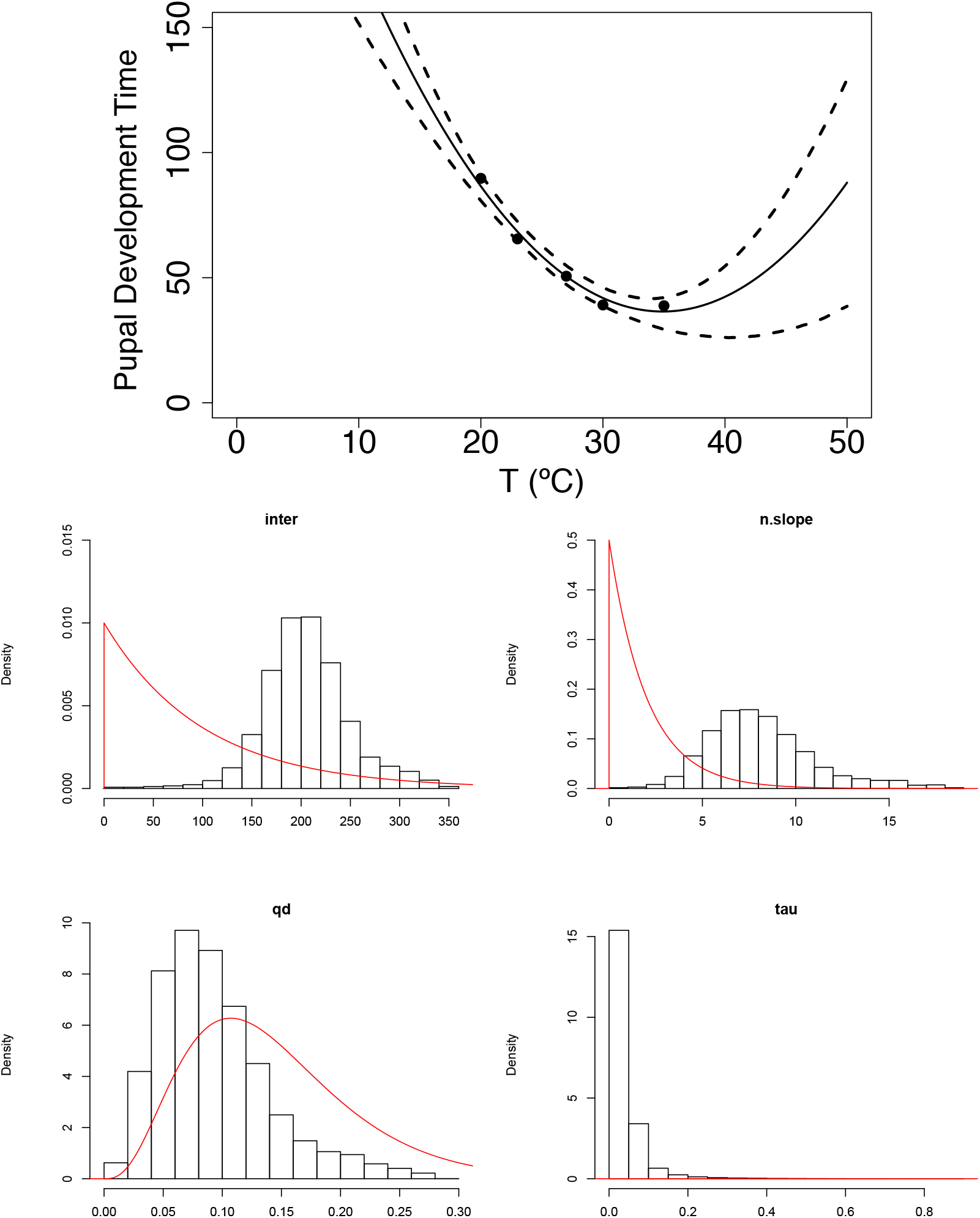
(Top) The mean trajectory in solid line and HPD interval in dashed black for pupal development time *ρ_P_*. (Bottom) Histograms of the posterior distribution for each parameter of the quadratic fit for pupal development time *ρ_P_*. The prior distribution for each parameter is plotted in red. The quadratic fit is determined by the equation *inter – n.slope T* + *qd T*^2^ using a normal distribution with precision *τ*.

#### Thermal traits prior distributions

Table S1.2 summarizes all the priors used to fit the thermal curves.

**Figure S1.11:**
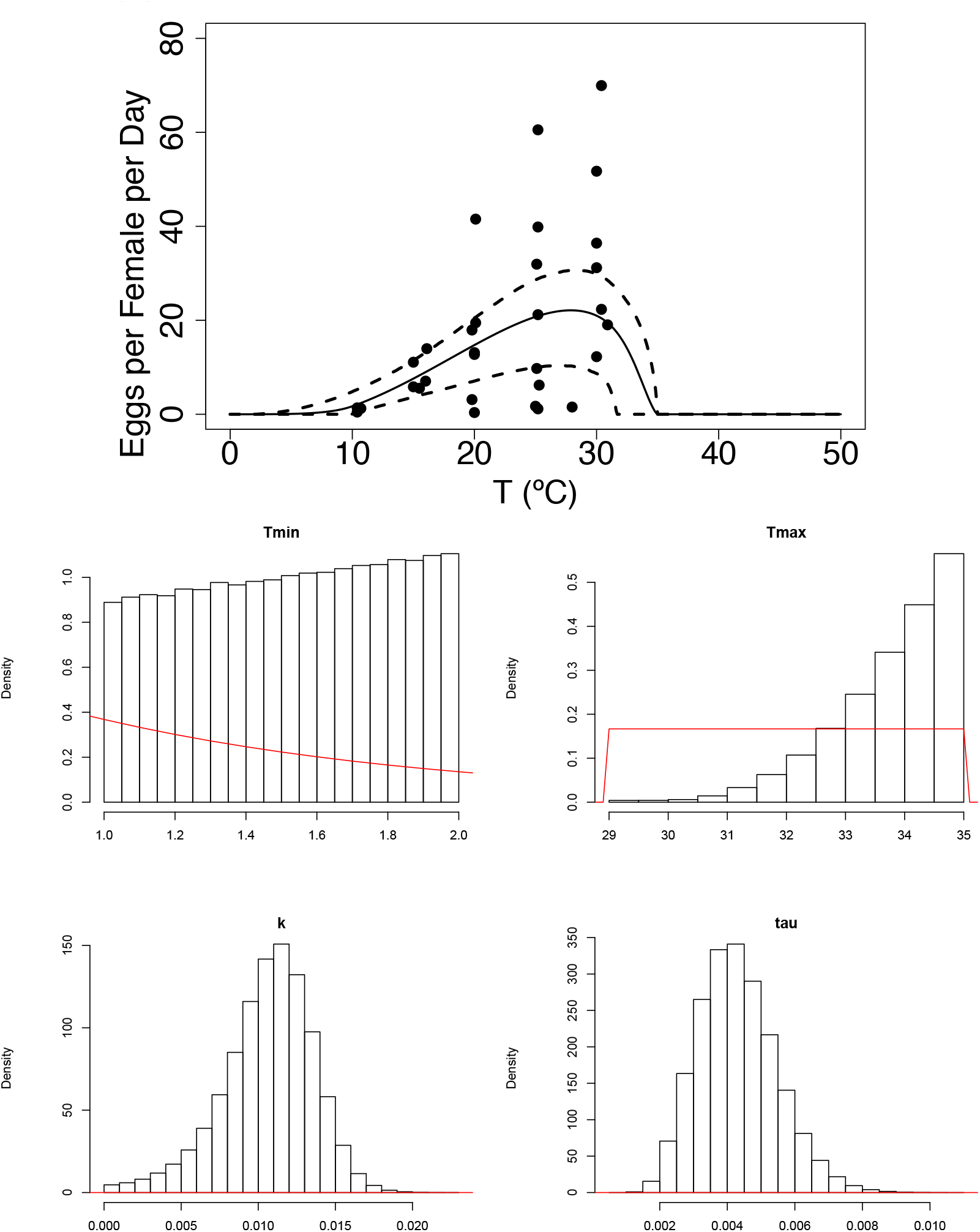
(Top) The mean trajectory in solid line and HPD interval in dashed black for fecundity *F*. (Bottom) Histograms of the posterior distribution for each parameter of the Brière fit for fecundity *F*. The prior distribution for each parameter is plotted in red. The Brière fit is determined by the equation 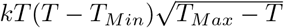 using a normal distribution with precision *τ*.

### S1.5 Posterior distributions for all *R*_0_ forms

For all three *R*_0_ posterior distributions we provide posterior distributions for the lower temperature limit, peak temperature, and upper temperature limit.

**Figure S1.12:**
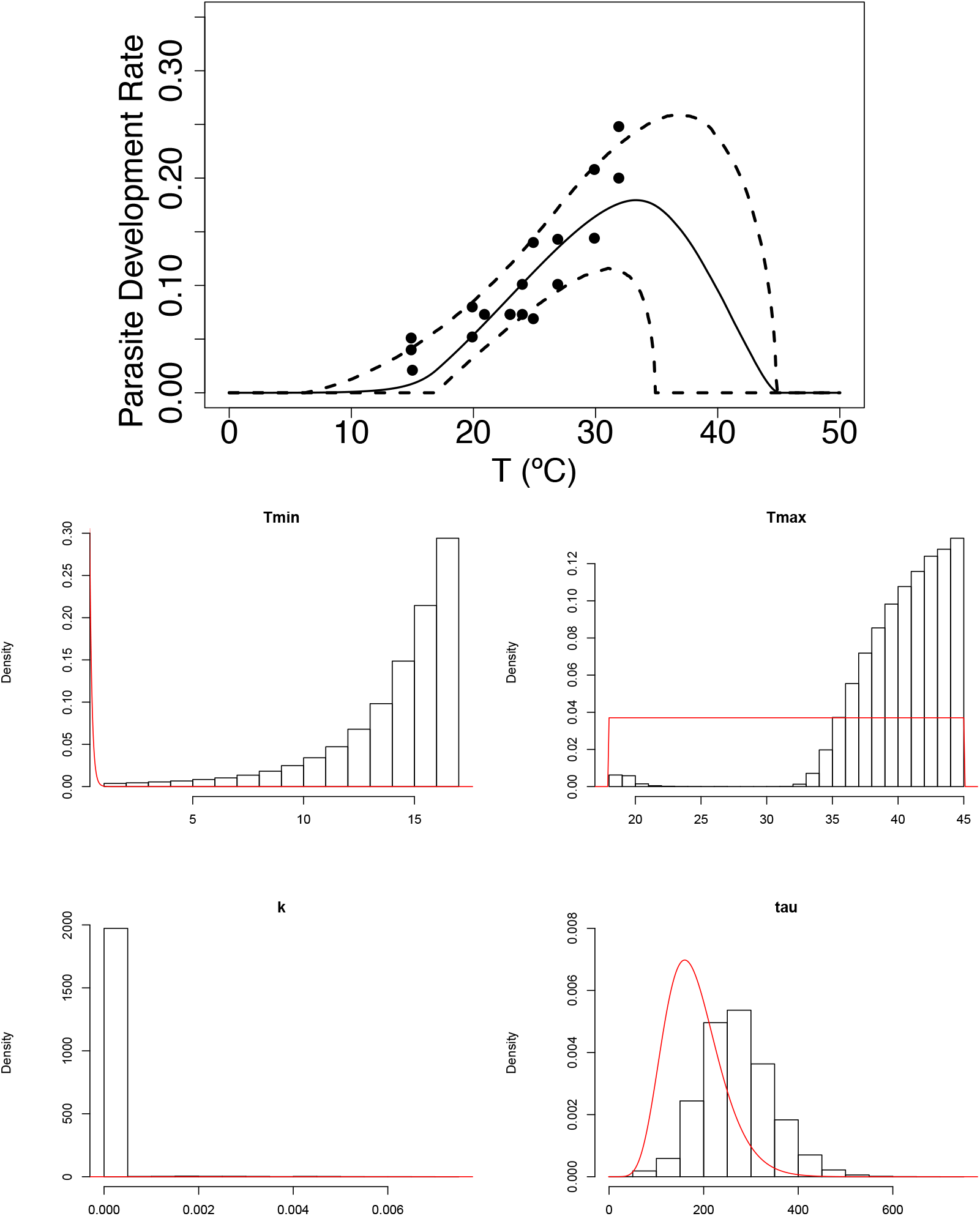
(Top) The mean trajectory in solid line and HPD interval in dashed black for the parasite development rate *ν*. (Bottom) Histograms of the posterior distribution for each parameter of the Brière fit for the parasite development rate *ν*. The prior distribution for each parameter is plotted in red. The Brière fit is determined by the equation 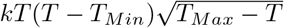 using a normal distribution with precision *τ*.

**Figure S1.13:**
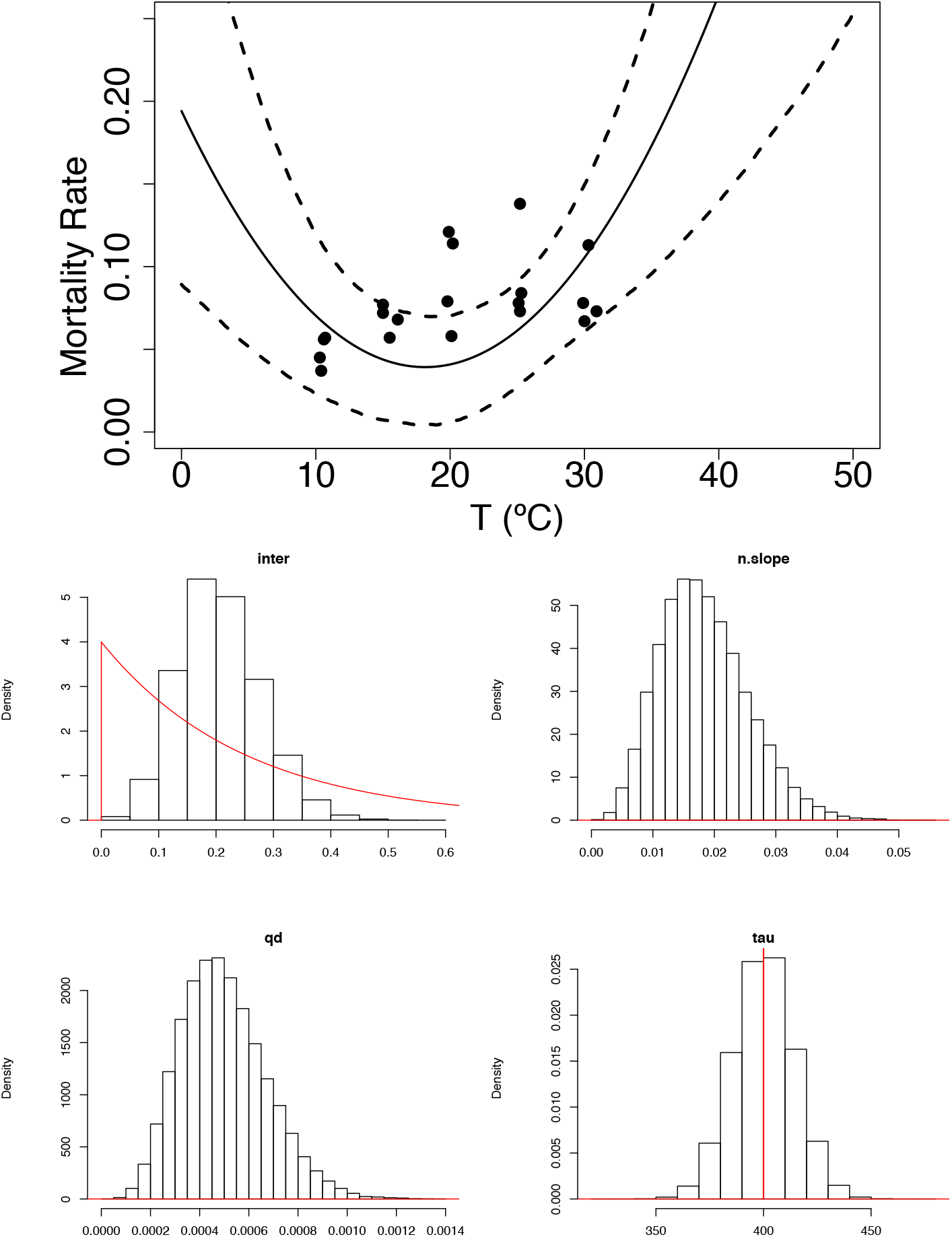
(Top) The mean trajectory in solid line and HPD interval in dashed black for the mortality rate *μ*. (Bottom) Histograms of the posterior distribution for each parameter of the quadratic fit for the mortality rate *μ*. The prior distribution for each parameter is plotted in red. The quadratic fit is determined by the equation *inter – n.slope T* + *qd T*^2^ using a normal distribution with precision *τ*.

**Table S1.2:** Prior distributions for each of the parameters for the fitting of the responses for each of the thermal traits considered.

| Model Parameter | Mean Function | Parameters | Prior |
| --- | --- | --- | --- |
| Biting Rate<br>$a$ | Brière | $T_{Min}$<br>$T_{Max}$<br>$k$<br>$\tau$ | dunif(0, 20)<br>dunif(20,40)<br>dgamma(1,20)<br>dgamma(0.01, 0.01) |
| Transmission probability<br>$b$ | Brière | $T_{Min}$<br>$T_{Max}$<br>$k$ | dunif(10,24)<br>dunif(25,35)<br>dgamma(1,10) |
| Egg Survival Probability<br>$p_E$ | Brière | $T_{Min}$<br>$T_{Max}$<br>$k$<br>$\tau$ | dunif(10,20)<br>dunif(35,40)<br>dgamma(1,20)<br>dgamma(7, $5^{-10}$) |
| Larval Survival Probability<br>$p_L$ | Brière | $T_{Min}$<br>$T_{Max}$<br>$k$<br>$\tau$ | dunif(0,8)<br>dunif(30,40)<br>dgamma(1,20)<br>dgamma(1.5, 0.001) |
| Pupal Survival Probability<br>$p_P$ | Brière | $T_{Min}$<br>$T_{Max}$<br>$k$<br>$\tau$ | dunif(1,5)<br>dunif(35,40)<br>dgamma(1,5)<br>dgamma(10, 0.002) |
| Egg Development Time<br>$\rho_E$ | Quadratic | inter<br>n.slope<br>qd<br>$\tau$ | dgamma(1, 0.01)<br>dgamma(1, 0.5)<br>dgamma(4,28)<br>dnorm(3, 1/800) |
| Larval Development Time<br>$\rho_L$ | Quadratic | inter<br>n.slope<br>qd<br>$\tau$ | dgamma(1, 0.01)<br>dgamma(1, 0.5)<br>dgamma(4,28)<br>dnorm(3, 1/1000) |
| Pupal Development Time<br>$\rho_P$ | Quadratic | inter<br>n.slope<br>qd<br>$\tau$ | dgamma(1, 0.01)<br>dgamma(1, 0.5)<br>dgamma(4,28)<br>dnorm(3, 1/200) |
| Eggs per Female per Day<br>$F$ | Brière | $T_{Min}$<br>$T_{Max}$<br>$k$<br>$\tau$ | dunif(1, 10)<br>dunif(29,35)<br>dgamma(1,1)<br>dgamma(9, 0.0005) |
| Parasite Development Rate<br>$\nu$ | Brière | $T_{Min}$<br>$T_{Max}$<br>$k$<br>$\tau$ | dunif(1, 17)<br>dunif(18,45)<br>dgamma(1,10)<br>dgamma(9, 0.05) |
| Adult Mortality Rate<br>$\mu$ | Quadratic | inter<br>n.slope<br>qd<br>$\tau$ | dgamma(2,2)<br>dgamma(3,3)<br>dgamma(2,2)<br>dnorm(1000, 1/500) |

**Figure S1.14:**
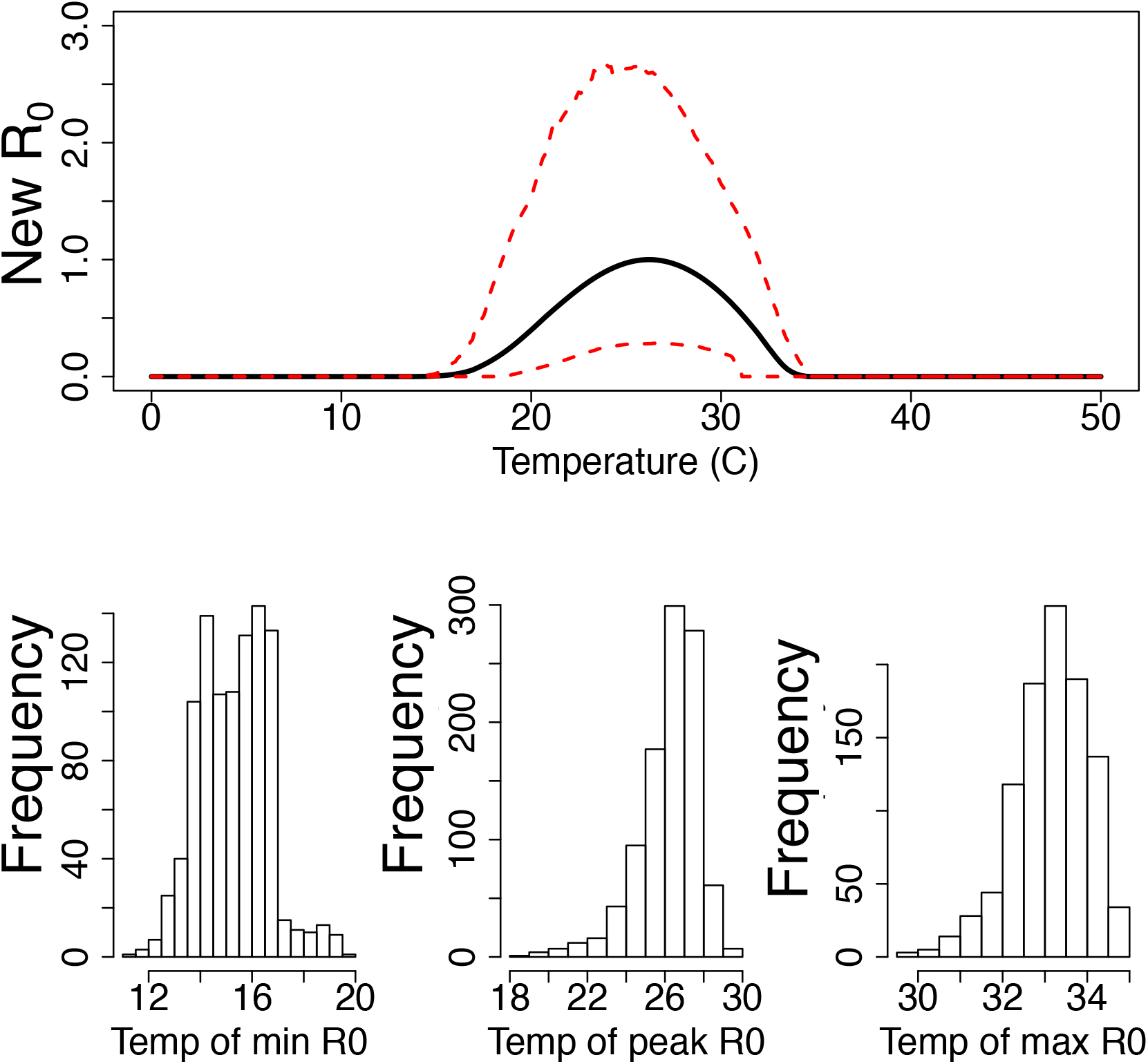
Minimum, peak and, maximum temepratures posterior densities for the *R*_0_ presented here.

**Figure S1.15:**
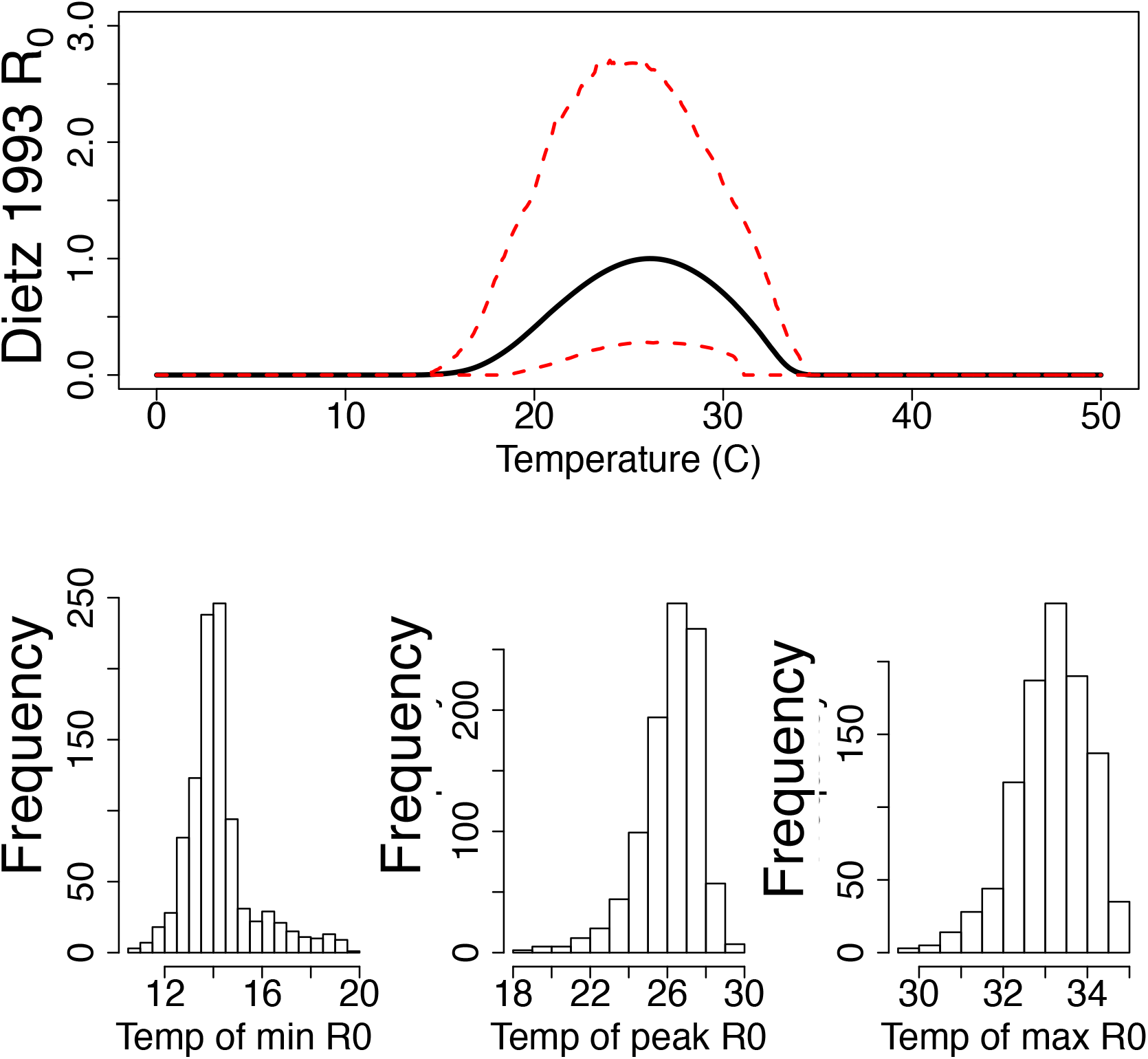
Minimum, peak and, maximum temepratures posterior densities for Dietz 1993 [24] *R*_0_

**Figure S1.16:**
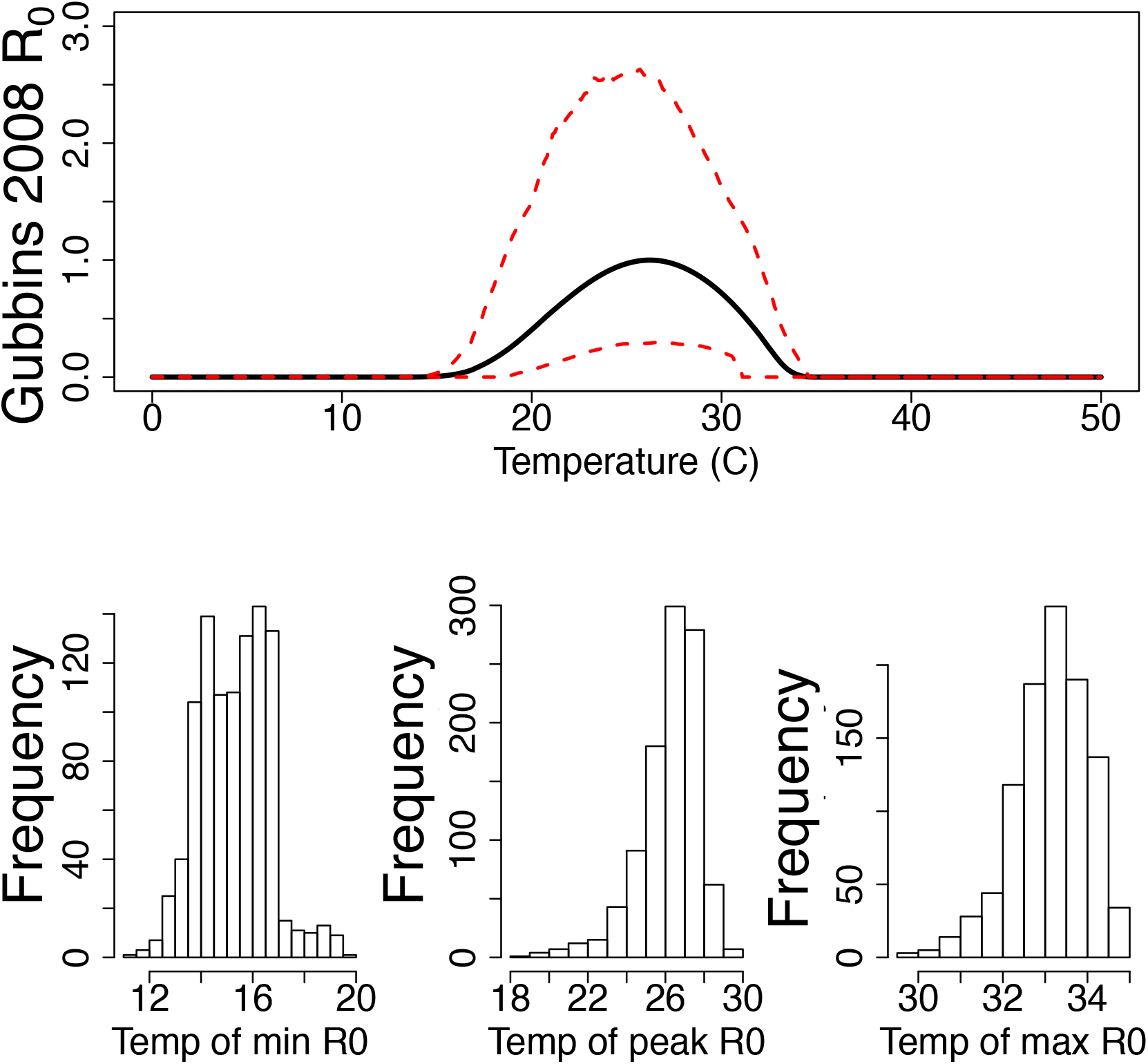
Minimum, peak and, maximum temepratures posterior densities for Gubbins 2008 [19] *R*_0_

## References

[1] E. A. Mordecai, K. P. Paaijmans, L. R. Johnson, C. Balzer, T. Ben-Horin, E. de Moor, A. McNally, S. Pawar, S. J. Ryan, T. C. Smith, et al., “Optimal temperature for malaria transmission is dramatically lower than previously predicted,” Ecology letters, vol. 16, no. 1, pp. 22–30, 2013.

[2] L. R. Johnson, T. Ben-Horin, K. D. Lafferty, A. McNally, E. Mordecai, K. P. Paaijmans, S. Pawar, and S. J. Ryan, “Understanding uncertainty in temperature effects on vector-borne disease: a Bayesian approach,” Ecology, vol. 96, no. 1, pp. 203–213, 2015.

[3] R. A. Taylor, E. A. Mordecai, C. A. Gilligan, J. R. Rohr, and L. R. Johnson, “Mathematical models are a powerful method to understand and control the spread of Huanglongbing,” PeerJ, vol. 4, p. e2642, 2016.

[4] C. Calisher and P. Mertens, “Taxonomy of African horse sickness viruses,” in African Horse Sickness, pp. 3–11, Springer, 1998.

[5] P. Mellor, J. Boorman, and M. Baylis, “Culicoides biting midges: their role as arbovirus vectors,” Annual review of entomology, vol. 45, no. 1, pp. 307–340, 2000.

[6] E. Wittmann, P. Mellor, and M. Baylis, “Effect of temperature on the transmission of orbiviruses by the biting midge, Culicoides sonorensis,” Medical and veterinary entomology, vol. 16, no. 2, pp. 147–156, 2002.

[7] S. Gubbins, S. Carpenter, M. Baylis, J. L. Wood, and P. S. Mellor, “Assessing the risk of bluetongue to UK livestock: uncertainty and sensitivity analyses of a temperature-dependent model for the basic reproduction number,” Journal of the Royal Society Interface, vol. 5, no. 20, pp. 363–371, 2007.

[8] W. J. Tabachnick, C. T. Smartt, and C. R. Connelly, “Bluetongue,” UF IFSAS Extension, 2008.

[9] A. S. Lear and R. J. Callan, Overview of Bluetongue, 2014.

[10] N. J. MacLachlan and A. J. Guthrie, “Re-emergence of bluetongue, African horse sickness, and other Orbivirus diseases,” Veterinary Research, 2010.

[11] N. Preparedness, A. Incident Coordination Center, Veterinary Services, and P. H. I. Service, Bluetongue standard operating procedure: an overview of etiology and ecology, 2016.

[12] K. A. Alexander, N. J. MacLachlan, P. W. Kat, C. House, S. J. O’Brien, N. W. Lerche, M. Sawyer, L. G. Frank, K. Holekamp, L. Smale, J. W. McNutt, M. K. Laurenson, M. G. L. Mills, and B. I. Osburn, “Evidence of natural bluetongue virus infection among african carnivores,” The American journal of tropical medicine and hygiene, 1994.

[13] W. O. for Animal Health, “Bluetongue,” World Organisation for Animal Health (OIE), 2013.

[14] M. Jenckel, E. Brèard, C. Schulz, C. Sailleau, C. Viarouge, B. Hoffmann, D. Höper, M. Beer, and S. Zientara, “Complete Coding Genome Sequence of Putative Novel Bluetongue Virus Serotype 27,” Microbiology Resource Announcements, vol. 3, no. 2, 2015.

[15] USDA, Veterinary Biological Products. United States Department of Agriculture, 2019.

[16] USDA, “U.S. Cattle & Beef Industry Statistics and Information,” Economic Research Service, 2015.

[17] USDA, “Orbiviruses Gap Analysis: Bluetongue and Epizootic Hemorrhagic Disease,” Agricultural Research Service, 2013.

[18] EC, “Bluetongue seasonally vector free periods,” European Comission, 2016.

[19] S. Gubbins, S. Carpenter, M. Baylis, J. L. Wood, and P. S. Mellor, “Assessing the risk of bluetongue to UK livestock: uncertainty and sensitivity analyses of a temperature-dependent model for the basic reproduction number,” Journal of the Royal Society Interface, vol. 5, no. 20, pp. 363–371, 2007.

[20] J. Turner, R. G. Bowers, and M. Baylis, “Two-host, two-vector basic reproduction ratio (R0) for bluetongue,” PloS one, vol. 8, no. 1, p. e53128, 2013.

[21] S. P. Brand, K. S. Rock, and M. J. Keeling, “The interaction between vector life history and short vector life in vector-borne disease transmission and control,” PLoS computational biology, vol. 12, no. 4, p. e1004837, 2016.

[22] R. Ross, The prevention of malaria. John Murray; London, 1911.

[23] G. Macdonald et al., “The epidemiology and control of malaria.,” The Epidemiology and Control of Malaria., 1957.

[24] K. Dietz, “The estimation of the basic reproduction number for infectious diseases,” Statistical methods in medical research, vol. 2, no. 1, pp. 23–41, 1993.

[25] O. Diekmann and J. A. P. Heesterbeek, Mathematical epidemiology of infectious diseases: model building, analysis and interpretation, vol. 5. John Wiley & Sons, 2000.

[26] O. Diekmann, J. A. P. Heesterbeek, and M. G. Roberts, “The construction of next-generation matrices for compartmental epidemic models,” Journal of the Royal Society Interface, p. rsif20090386, 2009.

[27] A. I. Dell, S. Pawar, and V. M. Savage, “Systematic variation in the temperature dependence of physiological and ecological traits,” Proceedings of the National Academy of Sciences, vol. 108, no. 26, pp. 10591–10596, 2011.

[28] M. J. Angilletta Jr and M. J. Angilletta, Thermal adaptation: a theoretical and empirical synthesis. Oxford University Press, 2009.

[29] W. J. Tabachnick, “Culicoides variipennis and bluetongue-virus epidemiology in the united states,” Annual review of entomology, vol. 41, no. 1, pp. 23–43, 1996.

[30] J. A. Huwaldt and S. Steinhorst, “Plot digitizer,” URL http://plotdigitizer.sourceforge.net, 2013.

[31] rjags: Bayesian graphical models using MCMC.

[32] L. Joseph, D. B. Wolfson, and R. D. Berger, “Sample size calculations for binomial proportions via highest posterior density intervals,” Journal of the Royal Statistical Society: Series D (The Statistician), vol. 44, no. 2, pp. 143–154, 1995.

[33] R: A Language and Environment for Statistical Computing.

[34] R. J. Hijmans, S. E. Cameron, J. L. Parra, P. G. Jones, and A. Jarvis, “Very high resolution interpolated climate surfaces for global land areas,” International Journal of Climatology: A Journal of the Royal Meteorological Society, vol. 25, no. 15, pp. 1965–1978, 2005.

[35] M. Gilbert, G. Nicolas, G. Cinardi, T. P. Van Boeckel, S. Vanwambeke, W. G. R. Wint, and T. P. Robinson, “Global sheep distribution in 2010 (5 minutes of arc),” 2018.

[36] M. Gilbert, G. Nicolas, G. Cinardi, T. P. Van Boeckel, S. O. Vanwambeke, G. W. Wint, and T. P. Robinson, “Global distribution data for cattle, buffaloes, horses, sheep, goats, pigs, chickens and ducks in 2010,” Scientific data, vol. 5, p. 180227, 2018.

[37] R. J. Hijmans, raster: Geographic Data Analysis and Modeling, 2019.

[38] R. J. Hijmans and J. van Etten, raster: Geographic analysis and modeling with raster data. R package version 2.0–12, 2012.

[39] R. Bivand, “Lewin-koh, n. maptools: Tools for reading and handling spatial objects. r packag. version 0.9-1. https,” CRAN. R-project. org/package= maptools, 2017.

[40] R. Bivand, T. Keitt, B. Rowlingson, E. Pebesma, M. Sumner, R. Hijmans, E. Rouault, and M. R. Bivand, “Package ?rgdal?,” Bindings for the Geospatial Data Abstraction Library. Available online: https://cran.r-project.org/web/packages/rgdal/index.html (accessed on 15 October 2017), 2015.

[41] S. J. Ryan, A. McNally, L. R. Johnson, E. A. Mordecai, T. Ben-Horin, K. Paaijmans, and K. D. Lafferty, “Mapping physiological suitability limits for malaria in africa under climate change,” Vector-Borne and Zoonotic Diseases, vol. 15, no. 12, pp. 718–725, 2015.

[42] N. Courtejoie, G. Zanella, and B. Durand, “Bluetongue transmission and control in Europe: A systematic review of compartmental mathematical models,” Preventive veterinary medicine, 2018.

[43] N. Hartemink, B. Purse, R. Meiswinkel, H. E. Brown, A. De Koeijer, A. Elbers, G.-J. Boender, D. Rogers, and J. Heesterbeek, “Mapping the basic reproduction number (R0) for vector-borne diseases: a case study on bluetongue virus,” Epidemics, vol. 1, no. 3, pp. 153–161, 2009.

[44] E. A. Mordecai, J. M. Cohen, M. V. Evans, P. Gudapati, L. R. Johnson, C. A. Lippi, K. Miazgowicz, C. C. Murdock, J. R. Rohr, S. J. Ryan, et al., “Detecting the impact of temperature on transmission of Zika, dengue, and chikungunya using mechanistic models,” PLoS neglected tropical diseases, vol. 11, no. 4, p. e0005568, 2017.

[45] R. A. Taylor, S. J. Ryan, C. A. Lippi, D. G. Hall, H. A. Narouei-Khandan, J. R. Rohr, and L. R. Johnson, “Predicting the fundamental thermal niche of crop pests and diseases in a changing world: a case study on citrus greening,” Journal of Applied Ecology, vol. 56, no. 8, pp. 2057–2068, 2019.

[46] A. E. Jones, J. Turner, C. Caminade, A. E. Heath, M. Wardeh, G. Kluiters, P. J. Diggle, A. P. Morse, and M. Baylis, “Bluetongue risk under future climates,” Nature Climate Change, vol. 9, no. 2, p. 153, 2019.

[47] A. L. Lloyd, “Realistic distributions of infectious periods in epidemic models: changing patterns of persistence and dynamics,” Theoretical Population Biology, vol. 60, no. 1, pp. 59–71, 2001.

[48] P. E. Parham and E. Michael, “Modeling the effects of weather and climate change on malaria transmission,” Environmental Health Perspectives, vol. 118, no. 5, p. 620, 2010.

[49] T. J. Lysyk and T. Danyk, “Effect of temperature on life history parameters of adult Culicoides sonorensis (Diptera: Ceratopogonidae) in relation to geographic origin and vectorial capacity for bluetongue virus,” Journal of Medical Entomology, vol. 44, no. 5, pp. 741–751, 2007.

[50] B. Mullens, A. Gerry, T. Lysyk, and E. Schmidtmann, “Environmental effects on vector competence and virogenesis of bluetongue virus in Culicoides: interpreting laboratory data in a field context,” Vet Ital, vol. 40, no. 3, pp. 160–166, 2004.

[51] S. Carpenter, A. Wilson, J. Barber, E. Veronesi, P. Mellor, G. Venter, and S. Gubbins, “Temperature dependence of the extrinsic incubation period of orbiviruses in Culicoides biting midges,” PloS one, vol. 6, no. 11, p. e27987, 2011.

[52] J. Vaughan and E. Turner Jr, “Development of immature Culicoides variipennis (Diptera: Ceratopogonidae) from Saltville, Virginia, at constant laboratory temperatures,” Journal of medical entomology, vol. 24, no. 3, pp. 390–395, 1987.

[53] C. D. Harvell, C. E. Mitchell, J. R. Ward, S. Altizer, A. P. Dobson, R. S. Ostfeld, and M. D. Samuel, “Climate warming and disease risks for terrestrial and marine biota,” Science, vol. 296, no. 5576, pp. 2158–2162, 2002.

